# Personalized functional networks scale with brain size beyond anatomy and are reorganized in 22q11.2 deletion syndrome

**DOI:** 10.64898/2026.11.16.767099

**Authors:** Kevin Y. Sun, William Snyder, Tien T. Tong, Marc Jaskir, Brooke L. Sevchik, Corey Horien, Benjamin Jung, Steven L. Meisler, Matthew Cieslak, Smrithi Prem, Eren Kafadar, Margaret Gardner, Justine Y. Hansen, Elizabeth Levitis, Katherine Cyr, Shivaram Karandikar, Jakob Seidlitz, Joëlle Bagautdinova, Kosha Ruparel, Hongming Li, Yong Fan, Russel T. Shinohara, Arielle S. Keller, Armin Raznahan, David R. Roalf, J. Eric Schmitt, Ruben C. Gur, Donna M. McDonald-McGinn, Raquel E. Gur, Theodore D. Satterthwaite, Aaron F. Alexander-Bloch

## Abstract

Anatomically, cortical regions scale differentially with total brain size: some show disproportionate contraction in smaller brains and expansion in larger brains, while others show relative stability. Whether personalized functional networks (PFNs) follow similar areal scaling principles, and how these principles shape functional reorganization in conditions that alter brain size, remain unknown. We investigated PFN scaling in typical development (ages 8-23; n=1,332) and 22q11.2 deletion syndrome (22q11DS), a neuropsychiatric genetic disorder with globally reduced brain size (62 cases, 153 matched controls; ages 10-24). In typical development, PFN scaling diverged from anatomy, deviating more from proportional scaling, particularly in association networks. In 22q11DS, accounting for expected scaling with brain size reduction revealed two robust network effects: relative peripheral visual reduction and relative cingulo-opercular enlargement. Vertex-level 22q11DS PFN reorganization beyond global cortical reduction showed spatial alignment with normative PFN changes associated with brain size, linking typical scaling principles to disorder-related alterations.

## Introduction

Total brain size varies almost twofold across people of the same age^1,2^. Cortical regions do not scale uniformly with this variation: some regions show disproportionate contraction in smaller brains and expansion in larger brains, while other regions show relative stability across different brain sizes^3,4^. Regional scaling differences are related to cortical evolution, development, cytoarchitecture, and functional brain networks^3^. While functional networks are typically defined at a group level, personalized functional networks (PFNs) can be mapped within each individual using functional MRI. PFNs vary substantially in the size, shape, and location of networks across individuals^5–7^, and this variation is associated with cognition and behavior^7–9^. However, it is unknown whether PFNs follow the scaling principles observed in anatomically defined regions. This knowledge gap is critical for understanding normative brain organization as well as conditions that affect both brain size and function, such as 22q11.2 deletion syndrome (22q11DS), a genetic disorder characterized by neuropsychiatric risk^10^ and global reductions in total cortical size^11–13^. Accounting for PFN scaling may disambiguate network-specific effects from expected consequences of a smaller brain.

Cortical scaling relationships are studied using allometry, a biological framework for modeling growth of a part relative to the whole^14^. In cortex, allometry distinguishes proportionate scaling (isometry) from supralinear scaling (hyperallometry), where regions change more than proportionately with total size, and sublinear scaling (hypoallometry), where regions change less than proportionately (**Fig. 1A**). Anatomical allometry studies of cortical surface have reported hierarchical patterns, with higher-order association networks tending towards hyperallometry and lower-order sensorimotor networks tending towards hypoallometry^3,4^. Applying group-level network boundaries to individual cortical surfaces captures how network territory scales with anatomy, but not individual differences in the spatial layout of networks, which are most pronounced in association cortex^5–7^. Although personalized network areas have been suggested to scale allometrically with total cortical surface area in adults^15^, it is unknown whether and how PFN allometry differs from purely anatomical allometry. In this study, our first aim was to characterize the allometry of PFNs in typical neurodevelopment.

**Figure 1.**
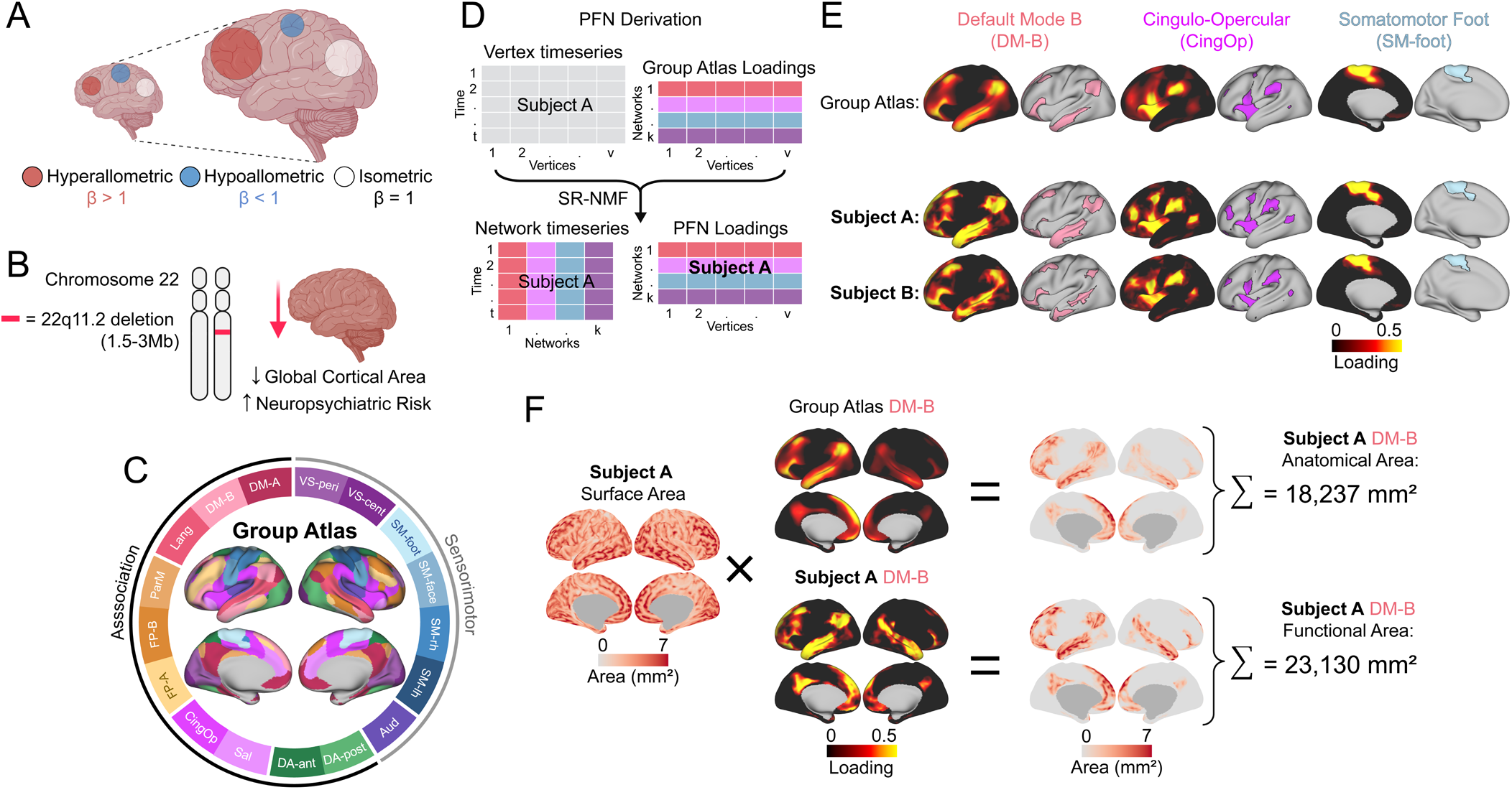
Study overview for the investigation of personalized functional network scaling and 22q11.2 deletion syndrome. **A.** Cortical area allometry defines how cortical areas scale proportionally with the entire cortical surface, with regions showing linear scaling (isometry), supralinear scaling (hyperallometry), or sublinear scaling (hypoallometry). **B**. 22q11.2 deletion syndrome (22q11DS) is a rare copy number variant disorder associated with both global reduction in cortical surface area and increased neuropsychiatric risk. **C.** Group atlas hard parcellation for reference (soft parcellation maps found in Supplementary Fig. 1). Sensorimotor and association designations notated by outer ring. **D.** Personalized functional network vertex-level loadings were derived from subject fMRI time series through regularized non-negative matrix factorization (NMF) using a consensus group atlas as a prior. t = 16:54 minutes, k = 17 networks, v = 59,412 vertices. **E.** Select group atlas network maps, soft-parcellated (loadings mapped) and hard-parcellated (winner-take-all), along with corresponding network maps from two exemplar subjects. **F.** Two network surface area measures were calculated across subjects using subject-specific vertex surface areas: anatomical area based on group-atlas-defined topography loadings and functional area based on PFN-defined topography loadings. Both measures were then associated with total cortical surface area, modeled allometrically (A), and 22q11.2 deletion syndrome (22q11DS) genotype (B). **Network abbreviations**: Aud = auditory; CingOp = cingulo-opercular; DA-ant = anterior dorsal attention; DA-post = posterior dorsal attention; DM = default mode; FP = frontoparietal; Lang = language; ParM = parietal memory; Sal = salience; SM = somatomotor; SM-lh = left hand; SM-rh = right hand; VS-cent = central visual; VS-peri = peripheral visual.

The importance of understanding PFN allometry is illustrated by functional brain organization in 22q11DS, a rare copy number variant disorder with elevated risk of psychosis^10^, cognitive deficits^16^, and global cortical surface area reduction^11–13^ (**Fig. 1B**). Regional surface area alterations in 22q11DS are well characterized, showing greater reductions in posterior compared to anterior regions^12,17,18^. Functional network dysconnectivity in 22q11DS has also been described^19^. Furthermore, outside of rare genetic variants, PFN topography is associated with psychiatric outcomes and its genetic risk^20–22^. However, whether the size, shape, and location of PFNs differ in 22q11DS remains unknown. A decrease in global cortical size is expected to produce reductions of varying magnitudes across networks, depending on how each network scales with cortical size. Differentiating network-specific effects from these expected size-related differences requires accounting for allometric scaling, and could identify candidate network substrates of 22q11DS psychiatric risk. Thus, the second aim of this study was to determine whether PFN differences in 22q11DS persist above and beyond global surface area reduction.

For our first aim, we investigated typical functional network allometry using data from the Philadelphia Neurodevelopmental Cohort (PNC)^23^, a community-based study of brain development (n=1,332, ages 8-23, 54% female). Because association cortex is less tightly coupled to underlying cortical structure than sensorimotor cortex^24–26^, displaying greater inter-individual variability in functional organization^7,27^, we hypothesized that PFN scaling would diverge from anatomical scaling and show greater deviation from isometry, particularly in association networks. We further asked whether this deviation would amplify the sensorimotor-association (S-A) pattern observed in anatomical allometry^3^ or depart from this hierarchical pattern. For our second aim, we investigated 22q11DS using data acquired using the same scanner and sequences as PNC^28^, with PNC controls matched on age, sex, race, and head motion (see Methods; matched sample ages 10-24, 46% female: n=62 22q11DS cases and n=153 PNC controls). We hypothesized that 22q11DS would show network-specific alterations beyond those expected from global cortical reduction. Finally, because 22q11DS disrupts neurodevelopmental processes^29^ that also shape cortical scaling across individuals^3,30^, we examined whether 22q11DS network reorganization spatially corresponded with normative topography scaling.

## Results

PFNs were derived^31^ from concatenated resting-state and task fMRI data^32^ (see Methods), using a group atlas from past work in PNC as a prior^7^ (hard parcellation, **Fig. 1C**; soft parcellations, **Supplementary Fig. 1**). This process resulted in a PFN loading matrix, in which loadings define the extent to which each vertex belongs to each network, a probabilistic parcellation we refer to as PFN topography (**Fig. 1D-E**). Surface areas of networks were calculated in two ways (**Fig. 1F**): anatomical network area, based on group-atlas-defined networks, and functional network area, based on each individual’s PFN topography. Anatomical area varied across subjects only through their subject-specific vertex-wise surface areas, with network topography held fixed. In contrast, functional area varied through both subject-specific vertex-wise surface area and network topography. These two measures enabled comparison of functional and anatomical allometry. In both cases, allometry was operationalized using log-log regression of network area on total cortical surface area (TSA)^14^, where slope β indicates isometry (β=1), hyperallometry (β>1), or hypoallometry (β<1). Effects described as “allometric” refer to those that deviate significantly from isometry in either direction.

As detailed below, our results characterized functional allometric deviation from anatomy and identified PFN effects in 22q11DS. In typical development, functional allometric coefficients spanned β=0.82-1.29 (mean |β-1|=0.10), consistent with greater deviation from isometry compared to atlas-based anatomical scaling (β=0.93-1.07, mean |β-1|=0.03). Diverging from the S-A axis, association networks showed heterogeneous scaling as opposed to uniform hyperallometry. In 22q11DS, accounting for global cortical reduction and individual network topography increased specificity of network effects. Widespread unadjusted reductions narrowed to two scaling-adjusted PFN effects: a relative reduction of the peripheral visual network (−5.9%) and relative enlargement of the cingulo-opercular network (+4.3%). At the vertex level, 22q11DS was associated with topographical reorganization that showed spatial correspondence with topography changes associated with normative scaling.

### PFN-derived functional scaling in typical development deviates more from isometry compared to anatomical scaling

Consistent with our hypothesis, functional areal scaling showed greater deviations from isometry than anatomical areal scaling (**Fig. 2**). Log-log models revealed that compared to group atlas-derived anatomical scaling, which spanned β=0.93-1.07 (**Fig. 2A**), PFN-derived functional scaling spanned a larger range of β=0.82-1.29 (**Fig. 2B**) and exhibited a greater average deviation from isometry (mean |β−1|=0.10 vs 0.03; all log-log plots shown in **Supplementary Fig. 2**). Functional and anatomical scaling coefficients were moderately correlated across networks (*r*=0.57; **Fig. 2C**), indicating that functional scaling partly reflects anatomical scaling while departing substantially from it: of seven allometric PFNs, six showed significantly different (*p_Bonf_*<0.05) scaling effects in PFN-defined versus group-atlas-defined networks (**Fig. 2C**) and five showed significantly greater deviations from isometry (*p_Bonf_*<0.05; **Supplementary Table 1**). In summary, accounting for inter-individual differences in functional network topography revealed stronger nonlinear scaling effects.

**Figure 2.**
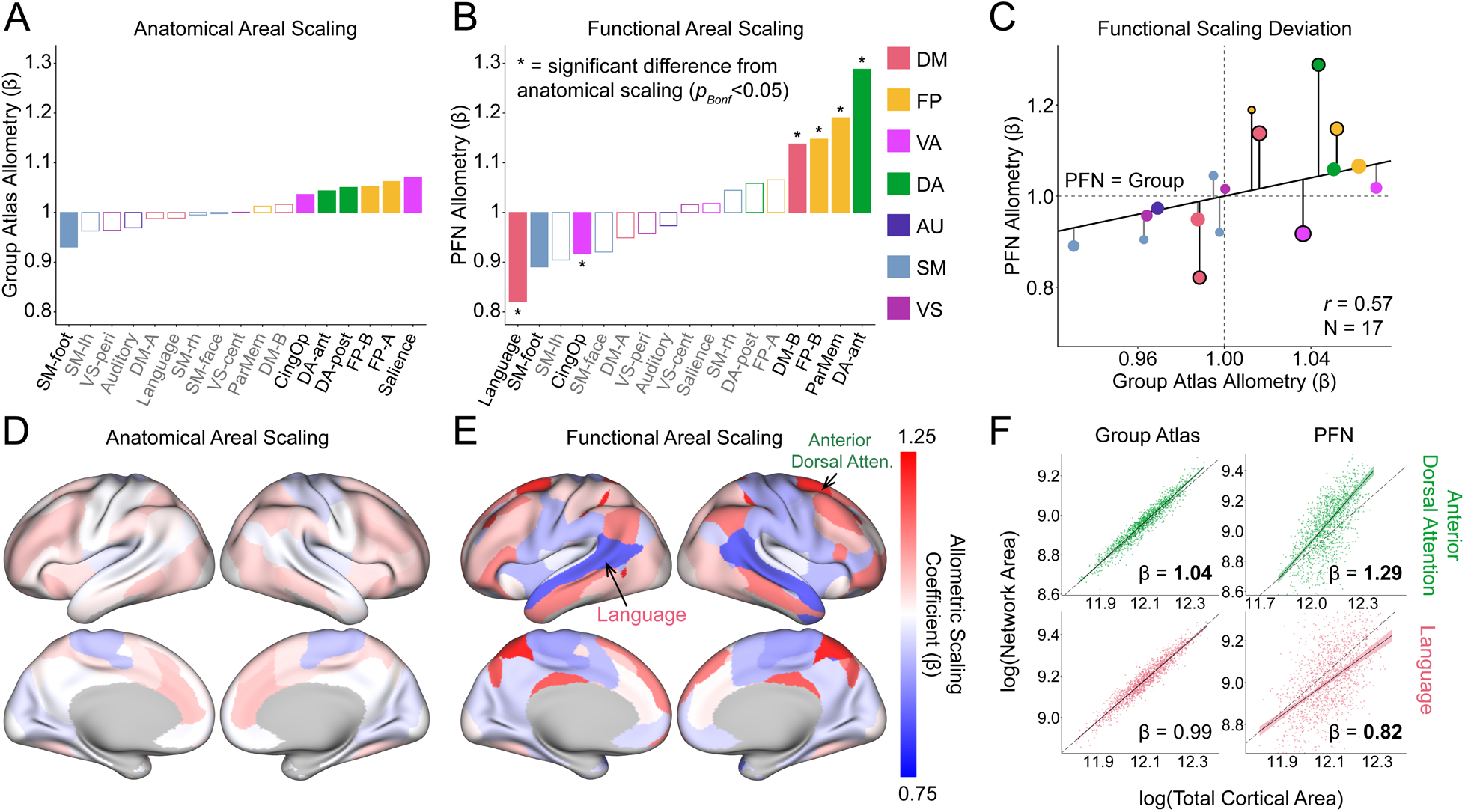
PFN-defined functional scaling in typical development revealed greater deviations from isometry compared to anatomical scaling. Bar charts showing anatomical areal scaling as defined by group atlas networks (**A**) and functional areal scaling defined by personalized functional networks (**B**), with significant deviation from isometry indicated by filled-in bars (p_Bonf_ < 0.05) and asterisks indicating significant deviation (p_Bonf_ < 0.05) of functional from anatomical areal scaling as assessed by two-sided p-values derived from inversion of bias-corrected and accelerated (BCa) bootstrap confidence intervals (5,000 resamples, see Methods). **C.** Functional areal scaling deviation from anatomical areal scaling, as shown by a scatterplot of PFN and group atlas allometry coefficients, showing an equality line (PFN = Group) and deviations from that line. Black outlines indicate significant difference between group atlas and PFN allometry coefficients; size of points reflect group atlas network size. Spatial pattern of unthresholded anatomical (**D**) and functional (**E**) areal scaling. **F**. Representative log-log plots of group atlas and PFN scaling of language and anterior dorsal attention networks. Dotted line indicates isometry (β = 1), bolded β value indicates significant deviation from isometry (p_Bonf_ < 0.05). **Functional systems**: DM = default mode; FP = frontoparietal; VA = ventral attention; DA = dorsal attention; AU = auditory; SM = somatomotor; VS = visual. **Network abbreviations**: CingOp = cingulo-opercular; DA-ant = anterior dorsal attention; DA-post = posterior dorsal attention; ParMem = parietal memory; SM-lh = left hand; SM-rh = right hand; VS-cent = central visual; VS-peri = peripheral visual.

Although normative anatomical scaling revealed a pattern aligned with the S-A axis (**Fig. 2A**), functional areal scaling diverged from this pattern. Significant deviation between functional and anatomical areal scaling occurred exclusively in association networks (**Fig. 2C**), with those six networks deviating in both hyperallometric (anterior dorsal attention, parietal memory, frontoparietal B, and default mode B) and hypoallometric directions (language and cingulo-opercular). Sensorimotor networks were comparatively isometric, with only one network showing significant deviation from isometry (foot somatomotor, hypoallometry). Functional deviation from anatomical allometry was greater in association networks (|Δβ| mean±SD=0.10±0.08) compared to sensorimotor networks (|Δβ| mean±SD=0.04±0.03), consistent with heterogeneous functional divergence in the former and relative anatomical correspondence in the latter. Greater effect sizes in functional allometry compared to anatomical allometry, and their divergence from the S-A axis, are apparent in network-level brain maps (**Fig. 2D-E**). Illustrative log-log plots of the strongest hypoallometric (language) and hyperallometric (anterior dorsal attention) networks identified in Fig. 2B are displayed in **Fig. 2F**.

One of six networks that deviated from anatomical scaling, the cingulo-opercular network, showed a reversal of directionality from significant anatomical hyperallometry, β=1.04, to significant functional hypoallometry, β=0.92 (**Fig. 2A-B**; outlined pink data point in bottom right quadrant, **Fig. 2C**). In contrast, and representative of reduced functional-anatomical deviation in sensorimotor networks overall, the foot somatomotor network showed no significant difference between functional (β=0.89) and anatomical (β=0.93) allometry (*p_Bonf_*=1.00; blue leftmost data point, **Fig. 2C**). This suggests that the sole example of significant sensorimotor network allometry is substantially attributable to anatomical variation, unlike the six allometric association networks showing significant functional-anatomical deviation.

A number of sensitivity analyses were implemented. As PNC spans ages 8-23, to ensure estimates were not influenced by age-related covariation, we repeated our analyses using models without an age spline covariate term, resulting in highly consistent results (**Supplementary Fig. 3**). Additionally, we tested consistency of our functional areal scaling results with previously published PFN work in the Human Connectome Project^15^, based on cross-atlas homologous network pairs between the two studies identified by the Network Correspondence Toolbox^33^ (**Supplementary Fig. 4**, Spearman *ρ=*0.68, one-sided *p=*0.01, n=11 corresponding networks).

### Vertex-level PFN topography varies systematically with cortical size in typical development

To examine how PFN topography changes with cortical size at finer spatial scales, mass univariate models regressed each vertex’s network loading on log(TSA). Because loadings quantify degree of network membership rather than area, these models are linear-log rather than log-log: each coefficient reflects the change in a vertex’s loading per unit increase in log(TSA); positive or negative coefficients indicate that a vertex belongs more or less strongly to the network when total cortical size increases. Across all variable PFN loadings (i.e., any vertex network loading that varies across individuals), 30.3% (71,277) showed significant associations with TSA (**Extended Data Table 1**).

Focusing on the seven allometric networks identified above (**Fig. 2B**), we found widespread significant associations (*p_FDR_*<0.05) across frontal, parietal, and temporal cortex (**Fig. 3A**; all network significant regions shown in **Supplementary Fig. 5**). Across allometric association networks, hypoallometric networks contracted into core regions with increasing cortical size (e.g., language network into left posterior superior temporal cortex; cingulo-opercular network into the anterior insula; **Fig. 3A bottom**). In contrast, hyperallometric networks expanded beyond their canonical territory (e.g., anterior dorsal attention network into inferior parietal cortex; **Fig. 3A bottom**). All 17 network effect size maps are shown in **Supplementary Fig. 6**.

**Figure 3.**
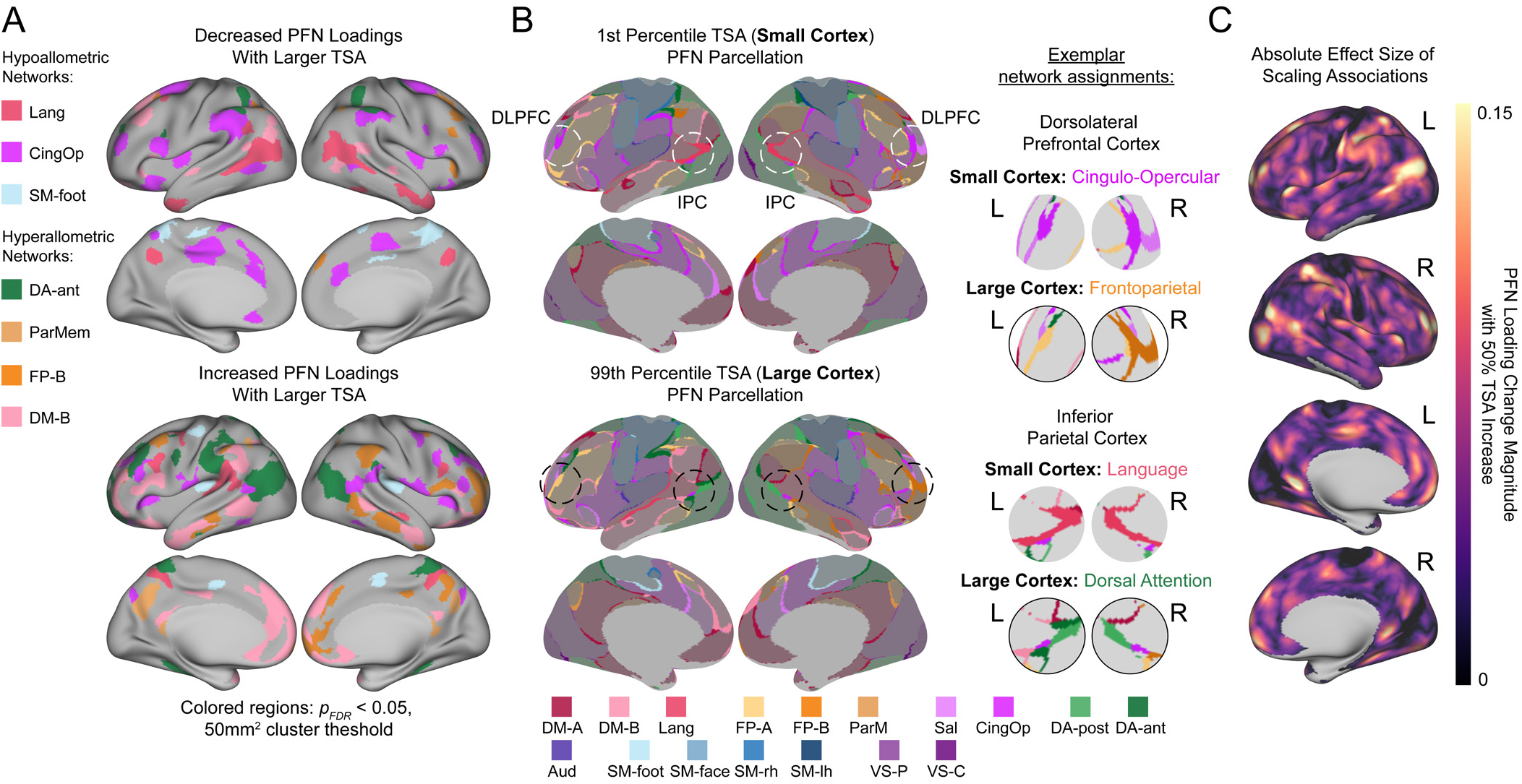
Vertex-level PFN scaling models in typical development. **A.** Significant (*p_FDR_* < 0.05) negative scaling effects (i.e., associated with cortical territory loss) and positive scaling effects (i.e., associated with cortical territory gain) of networks found to be hypo- or hyperallometric (as shown in Fig. 2B). 50mm^2^ cluster threshold applied; in cases of overlapping network vertices, the network with the larger absolute effect size is displayed. **B.** Hard-parcellated maps based on vertex-level model predictions, reflecting 1^st^ percentile (**Small Cortex**) of total cortical surface area (TSA) and 99^th^ percentile (**Large Cortex**) of TSA. White and black circles highlight exemplar network assignments. Consistent network identity vertices are faded in full cortex view and white in magnified circles. **C.** Map of the maximum absolute scaling effect size across networks, visualized as the change in PFN loading magnitude associated with a 50% increase in TSA. **Network abbreviations**: Aud = auditory; CingOp = cingulo-opercular; DA-ant = anterior dorsal attention; DA-post = posterior dorsal attention; DM = default mode; FP = frontoparietal; Lang = language; ParMem/ParM = parietal memory; Sal = salience; SM = somatomotor; SM-lh = left hand; SM-rh = right hand; VS-C = central visual; VS-P = peripheral visual. **Regions**: DLPFC = dorsolateral prefrontal cortex; IPC = inferior parietal cortex.

Directionality of vertex-level scaling associations mirrored network-level functional allometry (**Fig. 3A**). Specifically, negative associations dominated in language (mean effect size of significant vertices, β=−0.058) and cingulo-opercular (mean β=−0.057) networks, reflecting decreased topographical representation in larger cortex. In contrast, positive associations dominated in anterior dorsal attention (mean β=0.082), parietal memory (mean β=0.089), frontoparietal B (mean β=0.046), and default mode B (mean β=0.052) networks, reflecting increased topographical representation in larger cortex (**Extended Data Table 1**). The foot somatomotor network showed weak averaged directional association (mean β=−0.013) and more balanced topography associations (**Fig. 3A**), consistent with functional areal hypoallometry driven more by morphometry than by network topography. Models testing the scaling of each network’s functional extent, which aggregated each network’s PFN loadings across template surface area, revealed consistent results, with significant scaling effects in the same association networks with the exception of frontoparietal B (**Supplementary Table 1**).

To better understand how functional network topographies manifest across different cortical sizes, we derived winner-take-all network parcellations based on model-predicted PFN loadings for a small cortex (1^st^ percentile of TSA: ~0.14 m^2^) and a large cortex (99^th^ percentile of TSA: ~0.22 m^2^) (**Fig. 3B**). Hard parcellations revealed differential network assignments with increasing cortical size. For example, language network was reassigned to dorsal attention networks in inferior parietal cortex, and cingulo-opercular network was reassigned to frontoparietal networks in dorsolateral prefrontal cortex (**Fig. 3B**). We additionally visualized the absolute magnitude of network topography changes with scaling, which showed that inferior parietal and dorsolateral prefrontal cortex displayed strong effect sizes relative to other cortical regions (**Fig. 3C**). Topographical tradeoffs were also characterized quantitatively and summarized in a weighted flow diagram (**Extended Data Fig. 1**), revealing territorial tradeoffs between association networks across the cortical hierarchy (e.g., language to anterior dorsal attention, cingulo-opercular to default mode B). Network parcellation changes with cortical size were further visualized in a movie spanning the 1^st^ to 99^th^ percentile of TSA (**Supplementary Video 1**).

To complement vertex-level functional topography findings, we also tested vertex-level anatomical surface area allometry, revealing spatial correspondence to previously reported cortical surface area allometry^3^ (r=0.50, *p_spin_*=2×10^−4^; **Extended Data Fig. 2**).

### Scaling adjustment revealed two 22q11DS network effects robust to individual topography

Having characterized PFN scaling in normative development, we next sought to describe PFN effects in 22q11DS, with and without adjusting for scaling with TSA. Consistent with prior reports, TSA was decreased in 22q11DS (**Fig. 4A**; −4.7%, *p*=8.28×10^−5^). As expected given reductions in TSA, there were widespread negative effects in both group-atlas- and PFN-defined networks prior to adjustment for scaling (**Fig. 4B-C left**).

**Figure 4.**
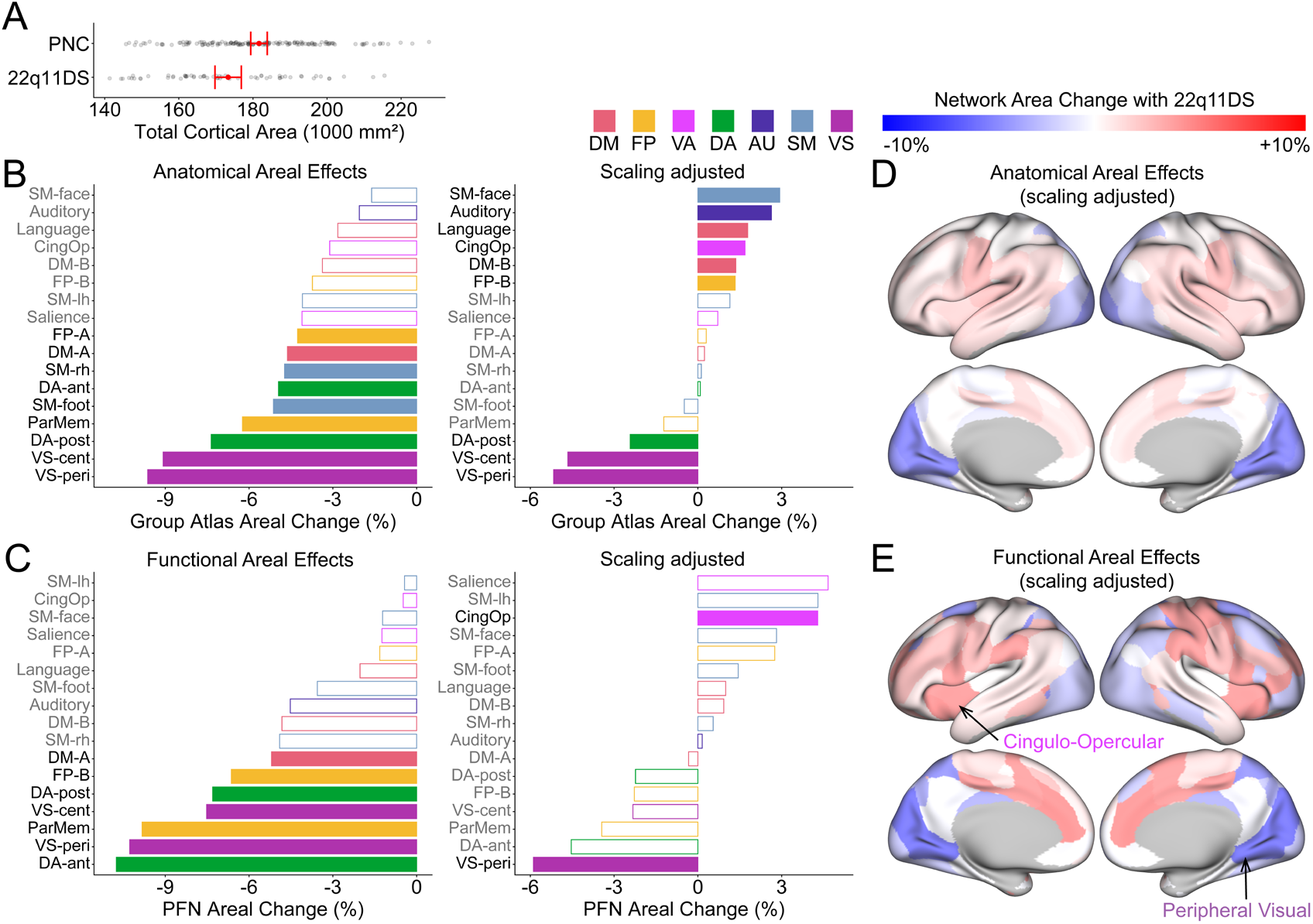
Surface area changes in 22q11DS functional networks after adjusting for scaling with total surface area. **A.** Effect of 22q11DS on total cortical surface area (raw data in grey, red shows covariate-adjusted model predictions and 95% confidence intervals). **B.** Anatomical areal effects of 22q11DS (i.e., effects on surface area of group atlas functional networks projected onto individual cortical surfaces) before and after adjusting for scaling. **C.** Functional areal effects of 22q11DS (i.e., effects on surface area of PFNs) before and after scaling adjustment; no significant differences were found between functional and anatomical effects (*p_Bonf_* ≥ 0.05). **D.** Spatial pattern of unthresholded scaling-adjusted anatomical areal effects of 22q11DS. **E.** Spatial pattern of unthresholded scaling-adjusted functional areal effects of 22q11DS, highlighting significant networks. **Functional systems**: DM = default mode; FP = frontoparietal; VA = ventral attention; DA = dorsal attention; AU = auditory; SM = somatomotor; VS = visual. The two significant effects identified aligned with the anterior-posterior axis: peripheral visual network in medial occipital cortex and cingulo-opercular network in anterior cingulate, anterior insula, and temporoparietal junction. **Network abbreviations**: CingOp = cingulo-opercular; DA-ant = anterior dorsal attention; DA-post = posterior dorsal attention; ParMem = parietal memory; SM-lh = left hand; SM-rh = right hand; VS-cent = central visual; VS-peri = peripheral visual.

Consistent with our hypothesis, we found network-specific effects beyond global cortical reduction, two of which remained significant when networks were individually defined (significant networks: scaling-unadjusted, 9 group atlas networks versus 7 PFN networks; scaling-adjusted, 9 group atlas networks versus 2 PFN networks). After scaling adjustment, group-atlas-derived models revealed anatomical areal differences relative to TSA, with six networks disproportionately larger and three disproportionately smaller (**Fig. 4B right**). Scaling-adjusted PFN-derived models revealed significance in a subset of these effects, with two network effects remaining: disproportionate reduction of the peripheral visual network (−5.9%, *p_Bonf_*=1.52×10^−4^) and disproportionate enlargement of the cingulo-opercular network (+4.3%, *p_Bonf_*=0.005) (**Fig. 4C right**; **Supplementary Table 2**).

Network-level 22q11DS areal results showed convergence with the anterior-posterior axis, a cortical gradient of neurogenesis^34^ and morphogenic patterning^35^ previously implicated in 22q11DS cortical surface area reductions^17,18^. Visualization of scaling-adjusted anatomical area effect sizes revealed consistent patterns: networks occupying occipital regions showed disproportionate relative reduction, whereas networks occupying frontal and temporal regions showed relative enlargement (**Fig. 4D**). This anterior-posterior organization was preserved in PFN-derived areal effects, although personalized network definitions displayed greater granularity within this overarching pattern (**Fig. 4E**).

We performed a number of sensitivity and exploratory analyses of network-level 22q11DS areal results. 22q11DS results were robust to age spline covariate exclusion (**Supplementary Fig. 7**). We next investigated if scaling-adjusted network developmental trajectories differed by 22q11DS status using age-by-genotype interaction models, and found that the cingulo-opercular network showed a significant interaction effect (*p_Bonf_*=0.030) in functional but not anatomical areas (**Extended Data Fig. 3A**). Specifically, relative enlargement of cingulo-opercular functional area in 22q11DS attenuated across adolescence (**Extended Data Fig. 3A**). Finally, in addition to adjusting for network scaling, we also tested if network scaling itself differed in 22q11DS versus normative development. We found a significant genotype-by-scaling interaction in frontoparietal network A (**Extended Data Fig. 3B**) using functional areas (*p_Bonf_*=0.020) but not anatomical areas (*p_Bonf_*=1.00), with 22q11DS cases showing a marked hypoallometry in functional scaling (β_22q_=0.62) relative to matched controls (β_PNC_=1.03). All interaction effects are shown in **Supplementary Table 3**.

### 22q11DS displayed vertex-level network reorganization beyond scaling adjustment

To examine PFN topography changes in 22q11DS at finer spatial scales, mass univariate models regressed each vertex’s network loading on genotype status, with and without log(TSA) as a covariate for scaling adjustment. Prior to scaling adjustment, 1.42% (3,335) of variable PFN vertex loadings showed significant associations with 22q11DS (*p_FDR_*<0.05, **Extended Data Table 2**). After scaling adjustment, 0.77% (1,808) of variable PFN vertex loadings showed significant associations with 22q11DS (*p_FDR_*<0.05, **Fig. 5A**, **Extended Data Table 2**). Although attenuated in number after adjustment, the spatial and directional pattern of significant loading changes remained broadly consistent across scaling-adjusted and unadjusted models (**Supplementary Fig. 8**).

**Figure 5.**
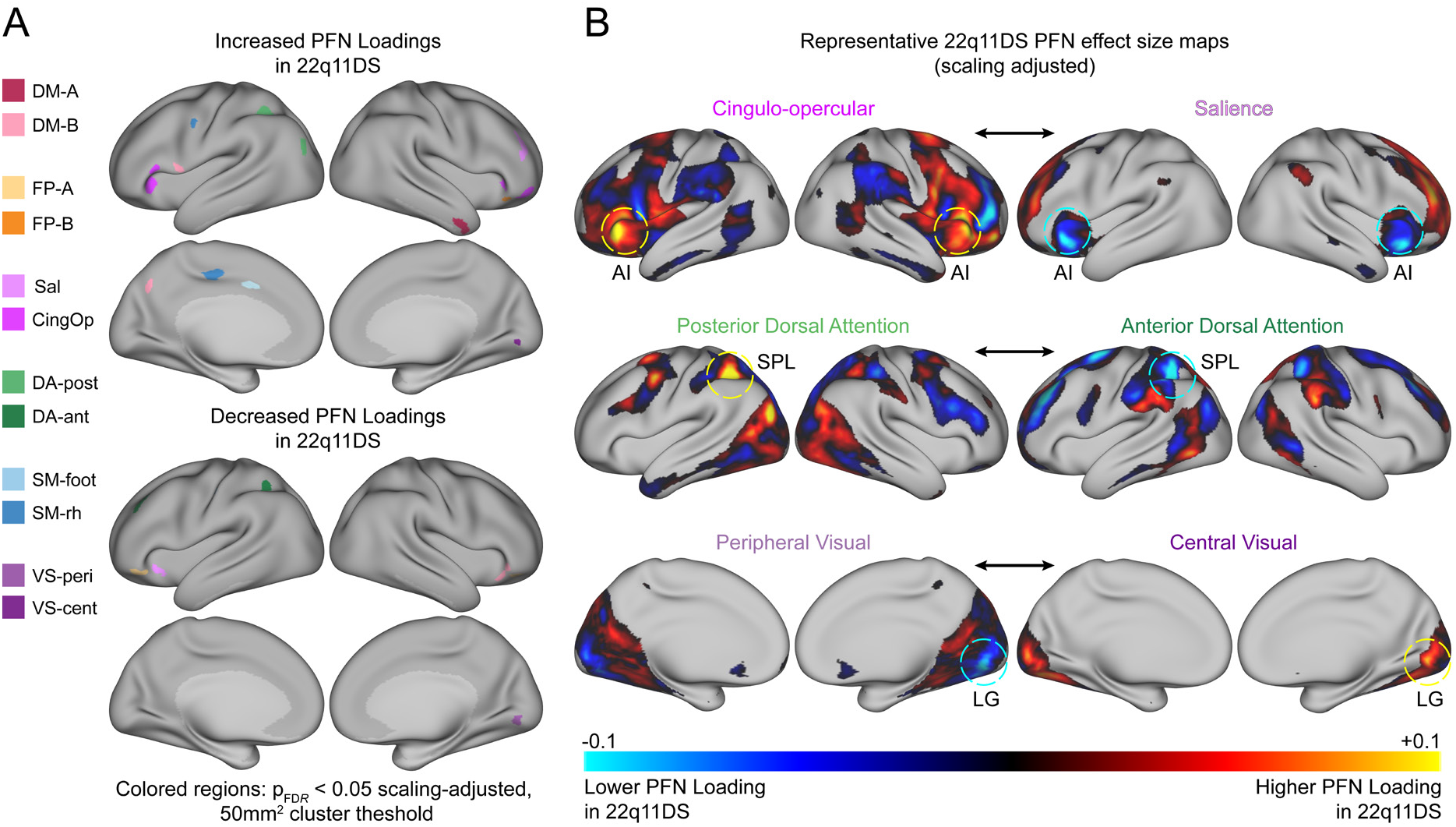
Vertex-level PFN 22q11DS models revealed distributed topography reorganization that persists beyond scaling adjustment. **A**. Significant (*p_FDR_* < 0.05) positive and negative 22q11DS network loading effects after scaling adjustment. 50mm^2^ cluster threshold applied; in cases of overlapping network vertices, the network with largest effect size is displayed. **B**. Representative unthresholded maps of 22q11DS vertex loading effects after scaling adjustment, showing functionally reciprocal network pairs (indicated by double-headed arrows). Yellow and blue dotted circles highlight regions with network loading tradeoffs. **Network abbreviations**: CingOp = cingulo-opercular; DA-ant = anterior dorsal attention; DA-post = posterior dorsal attention; DM = default mode; FP = frontoparietal; Sal = salience; SM = somatomotor; SM-rh = right hand; VS-cent = central visual; VS-peri = peripheral visual. **Regions**: AI = anterior insula; LG = lingual gyrus.

Vertex-level 22q11DS associations largely aligned with network-level functional area results (**Fig. 5**). Specifically, after scaling adjustment, significant negative associations were found in the right lingual gyrus of the peripheral visual network (mean β=−0.089), and significant positive associations were found across multiple regions of the cingulo-opercular network (mean β=0.093) (**Fig. 5A**; **Extended Data Table 2**). Additional significant loading associations were observed in default mode, frontoparietal, salience, dorsal attention, somatomotor, and visual networks (**Fig. 5A**).

Inspection of topography associations in 22q11DS revealed loading tradeoffs between networks (**Fig. 5B**; all scaling-adjusted 22q11DS effect size maps are shown in **Supplementary Fig. 9**). For example, decreased peripheral visual network loadings in the lingual gyrus were paired with increased central visual network loadings. Furthermore, the superior parietal lobule showed increased posterior dorsal attention network loadings but decreased anterior dorsal attention network loadings. Network tradeoffs were broadly consistent with and without scaling adjustment, but after scaling adjustment there was greater specificity in network tradeoffs, highlighting topography differences that were not attributable to reduction in TSA (**Extended Data Fig. 4**).

To complement vertex-level functional topography results, we investigated vertex-level surface area effects of 22q11DS, with and without scaling adjustment (**Extended Data Fig. 5**). Scaling adjustment revealed an anterior-posterior pattern of effects consistent with network-level 22q11DS results, with greater susceptibility to reduction in occipital regions and relative enlargement in insular, opercular, and temporal cortex.

### 22q11DS network reorganization overlaps with regions of normative size-related topography differences

Finally, we tested whether PFN topography effects of normative scaling and 22q11DS spatially corresponded. Effect size magnitude maps of normative scaling and scaling-adjusted 22q11DS were significantly correlated (*r=*0.34, *p_spin_*=2×10^−4^), indicating that cortical regions with greater PFN topography changes with TSA are also more likely to show topography changes in 22q11DS (**Fig. 6A**). Because regions with higher inter-individual variability in PFN topography, namely association cortex^7^, could show larger effects in both analyses, we residualized both effect-size maps for vertex-wise PFN variability (median absolute deviation across individuals, averaged across the 17 network loadings in PNC, n=1,332). Spatial correspondence between normative scaling and 22q11DS PFN effect sizes was attenuated but remained significant (*r=*0.17, *p_spin_*=2×10^−4^). In contrast, scaling-adjusted 22q11DS vertex-level surface area effects (**Extended Data Fig. 5**) showed no correspondence with normative anatomical allometry (**Extended Data Fig. 2**; *r=*0.01, *p_spin_*=0.93), indicating that topographical correspondence did not extend to regional surface area. We also tested for spatial correspondence between each PFN effect size map and eight maps of cortical hierarchy (**Supplementary Table 4**). Only the map of T1/T2 ratio, a proxy for intracortical myelination, was significantly correlated with the normative scaling effect map (*r*=−0.27, *p_Bonf_*=0.01), but did not survive residualization for PFN variability.

**Figure 6.**
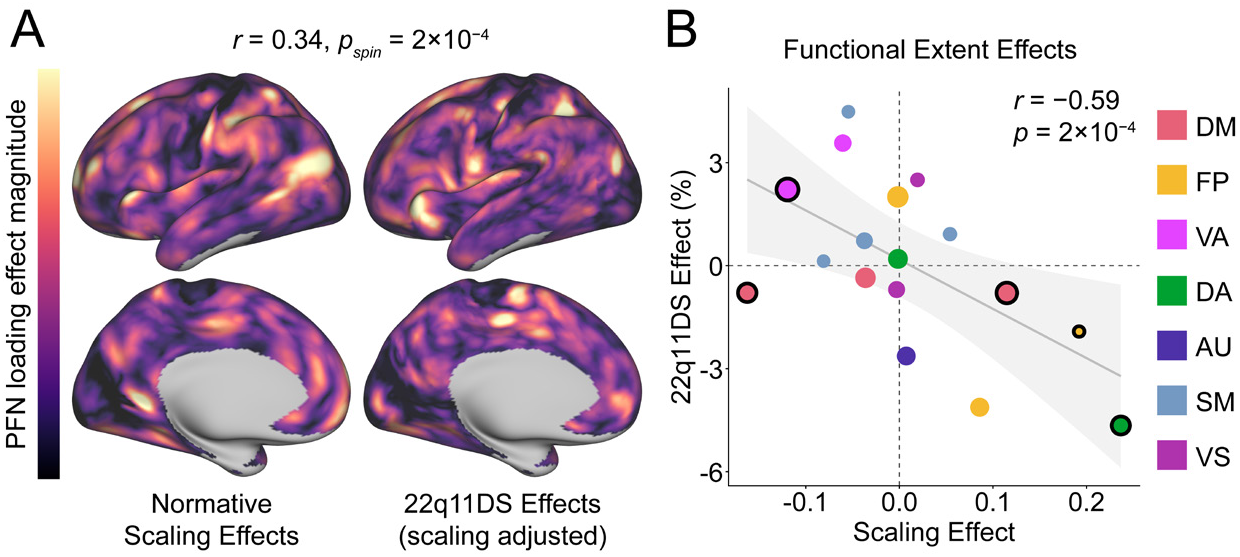
Shared cortical patterning underlies functional network topography differences in normative scaling and 22q11DS. **A.** Spatial correspondence was observed between effect size magnitude maps of PFN topography changes in normative scaling (also shown in Fig. 3C) and 22q11DS. After residualization of inter-individual PFN loading variability from both effect size maps, spatial correspondence attenuated but remained significant (*r*=0.17, *pspin*=2×10^−4^). **B.** Aggregating to the network-level, comparing the pattern of functional extent effects across scaling and 22q11DS after scaling adjustment revealed a significant negative correlation, assessed using two-sided p-values derived from inversion of bias-corrected and accelerated (BCa) bootstrap confidence intervals (5,000 resamples, see Methods). Black outlined networks correspond to significant functional extent scaling effects (Supplementary Table 1); size of points reflect group atlas network size. **Functional systems**: DM = default mode; FP = frontoparietal; VA = ventral attention; DA = dorsal attention; AU = auditory; SM = somatomotor; VS = visual.

Given this network-agnostic magnitude-based correspondence between normative scaling and 22q11DS effects, we next sought to understand how PFN loading effects related to each other in network and in direction. Aggregating to the network level, although no individual network’s functional extent differed significantly in 22q11DS after scaling adjustment (**Supplementary Table 2**), across networks, normative extent scaling was inversely correlated with scaling-adjusted 22q11DS extent effects (*r*=−0.59, 95% CI [−0.82, −0.42], *p*=2×10^−4^; **Fig. 6B**). Thus, networks with greater functional extent in smaller cortices normatively (analogous to hypoallometry, e.g., cingulo-opercular network) tended to be relatively expanded in 22q11DS after scaling adjustment, whereas networks with less functional extent in smaller cortices (analogous to hyperallometry, e.g., anterior dorsal attention network) tended to be relatively reduced. In other words, beyond its observed TSA reduction, 22q11DS network allocation shifted in the direction normatively associated with a smaller cortex.

## Discussion

In this study, we used an allometric approach to characterize how personalized functional network topography scales with cortical size variation in typical and 22q11DS neurodevelopment. In a community sample, functional network allometry differed markedly from anatomical allometry. Consistent with our hypothesis, personalized topography amplified deviations from isometry, but rather than aligning with the S-A axis, this amplification resulted in heterogeneous scaling across association networks. In youth with 22q11DS, adjusting for scaling and using personalized network definitions disentangled network-specific effects from widespread areal reductions, revealing two effects robust to inter-individual variation in topography. Finally, spatial alignment of functional scaling in the community sample with topographical alterations in 22q11DS suggests that shared organizational patterns underlie both typical functional network allometry and network reorganization in 22q11DS.

Prior anatomical allometry studies have described scaling aligned with the S-A axis, with association regions tending toward hyperallometry and sensorimotor regions toward hypoallometry^3,36^. Incorporating personalized network definitions revealed functional scaling that not only deviated more strongly from isometry, but also diverged from this S-A axis organization, with association networks spanning both hyper- and hypoallometry. The overall pattern of our network-level results broadly replicates a single prior report of functional network allometry in adults^15^. We extended this work to youth and, to our knowledge, provided the first comparison of functional network allometry with anatomical allometry, revealing and characterizing their divergence.

Functional divergence from anatomical scaling was concentrated in association networks, whereas sensorimotor networks more closely tracked anatomy. Higher-order association networks may be less tethered to early molecular signaling gradients and thalamic sensory input^37^, showing greater variability in functional organization^7,27^ as well as functional decoupling from intracortical microstructure^24^ and large-scale structural connectivity^25,26^. Thus, association networks may have greater flexibility for relative expansion *or* contraction with variation in total cortical size; in contrast, lower-order sensorimotor networks may be more tethered to these constraints^38^, remaining comparatively isometric. This cross-sectional pattern describes brain size covariation across individuals not explained by age, as results were consistent across age covariate specifications. Static allometry is nonetheless shaped by ontogeny: how networks scale across individuals of the same age partly reflects differences in their relative expansion trajectories earlier in development^30,39^. Indeed, our results are consistent with prior evidence that sensorimotor networks appear well-established in neonates while association networks display immature topography^40–42^, suggesting substantial association network individuation during the period of most rapid cortical growth^2,43^.

The direction of association network scaling was heterogenous. Across allometric association networks, hypoallometric networks appeared to contract toward their core regions with increasing cortical size^44,45^, whereas hyperallometric networks extended beyond their canonical territory^46^. More broadly, hypoallometric association networks were generally localized near paralimbic regions (i.e., transitional mesocortex including anterior insula, anterior cingulate, and temporal pole), whereas hyperallometric networks were distributed across heteromodal regions (i.e., six-layered isocortex: prefrontal cortex, temporoparietal junction, and precuneus)^47^. Prior anatomical allometry work reported a parallel cytoarchitectonic distinction, with hypoallometry enriched in limbic cortex (cingulate and parahippocampal cortex) and hyperallometry in association isocortex (lateral parietal, temporal and prefrontal cortex)^3^. Developmental literature suggests that, compared to heteromodal cortex, paralimbic regions show limited perinatal expansion^43,48^ and prolonged fetal sulcation^49–51^, as well as protracted developmental peaks in regional volume growth charts^2^ and lower relative areal expansion from infancy to adulthood^52^. These differential developmental trajectories are consistent with the heterogeneity observed across association network allometries.

Prior studies of functional networks in 22q11DS have relied on group-level network definitions; ours is among the first to map networks at the individual level. Given that 22q11DS is associated with globally reduced cortical surface area^12,13^, we modeled deletion effects on network area after accounting for scaling with total cortical size. These findings both converged with and refined the anterior-posterior organization of 22q11DS cortical reductions^17,18^. Specifically, PFN analysis isolated relative cortical reduction of posterior cortex to a single robust effect in the peripheral visual network, accompanied by loading tradeoffs in lingual gyrus, where peripheral visual loadings decreased and central visual loadings increased. Peripheral visual network reduction is consistent with altered functional activity of occipital regions reported previously in 22q11DS^53,54^, and may relate to deficits in face processing^28,55^ and visuospatial cognition^56^. PFNs also isolated relative 22q11DS cortical enlargement of anterior cortex to a single robust effect in the cingulo-opercular network, accompanied at the vertex level by increased cingulo-opercular loadings in anterior insula and frontal regions. This finding is consistent with prior reports of 22q11DS functional dysconnectivity^57,58^ in cingulo-opercular regions such as the insula, as well as relative enlargement of these regions in our own surface area models and in prior morphometric studies of 22q11DS^12,17,18^. Cingulo-opercular topographical reorganization may relate to psychosis vulnerability in 22q11DS, as prior literature has reported cingulo-opercular network abnormalities in the context of idiopathic psychosis^59–62^ and its genetic risk^22,63^.

Beyond the two network areal effects, personalized network topography in 22q11DS showed territorial reorganization across multiple networks not attributable to global cortical reduction. Scaling-adjusted 22q11DS topography effects were spatially correlated with normative scaling effects, a relationship that persisted after accounting for cortical patterning in inter-individual PFN variability^7^. Aggregating to the network level, normative scaling of functional extent was inversely correlated with scaling-adjusted 22q11DS extent effects: hyperallometric networks tended towards relative topographical reduction and hypoallometric networks tended towards relative expansion. That is, relative 22q11DS network allocation shifted in the direction normatively associated with a smaller cortex. These results suggest that 22q11DS may perturb developmental processes that coordinate network topography with cortical size, extending previous findings that neuropsychiatric alterations align with normative patterns of cortical organization and development^64,65^. In contrast, relative anatomical surface area differences in 22q11DS showed no spatial correspondence with normative anatomical allometry, consistent with recent multi-site findings^66^, suggesting that alignment of 22q11DS alterations with normative scaling may emerge specifically in functional network topography.

Network-specific differences in scaling and age-related patterns were found in 22q11DS relative to non-deleted youth. PFN area models revealed differential allometric scaling of frontoparietal network A depending on genotype: the network displayed hypoallometry in 22q11DS as opposed to near-isometry in non-deleted youth. Notably, altered allometric scaling has not been robustly observed in idiopathic autism^67^, raising the possibility that altered scaling is more characteristic of large-effect neurogenetic perturbation than of neurodevelopmental diagnosis generally. Furthermore, we observed age-related differences in 22q11DS: cingulo-opercular functional area showed a distinct developmental pattern in individuals with the deletion, with relative enlargement appearing to attenuate across adolescence. This parallels a prior report of an age-dependent transition from hyper-to hypoconnectivity in opercular cortex across puberty in 22q11DS^68^ and, more broadly, overlaps with the adolescent emergence of psychosis risk^69,70^.

The finding that scaling adjustment in PFN models of 22q11DS narrowed widespread reductions to two network-specific effects has implications beyond 22q11DS. Functional areal scaling diverged markedly from anatomical scaling, showing that cortical allometry is both network-specific and measure-dependent. Functional network studies of other sources of variation associated with brain size, such as sex^71–73^, cognition^74^, and other copy number variants^13^, as well as psychiatric disorders including autism^67^ and schizophrenia^75^, should consider personalized network definitions and allometric brain size adjustment.

Our results should be interpreted in the context of study limitations. First, by precision functional mapping standards, our concatenated fMRI scan time is limited at just under 17 minutes; however, recent work has shown high split-half reliability of PFN topography with as little as 10 minutes of data^9,76^. Second, although substantial given the deletion’s rarity^10^, our 22q11DS sample is small by statistical standards, including only 62 deletion cases; however, detection of group differences is supported by large prior effects of 22q11DS on structural and functional phenotypes^19,77^. Relatedly, this study did not investigate clinical or behavioral correlates within the 22q11DS cohort, which we would be underpowered to detect reliably^78^. Third, our analyses are cross-sectional, spanning late childhood to early adulthood, and thus developmental inferences, including links to early life patterning and age-related changes in 22q11DS, should be interpreted cautiously.

In summary, personalized functional networks revealed divergence of functional network allometry from anatomical scaling, identified 22q11DS network alterations beyond global cortical reduction that were robust to individual topography, and exhibited shared organizational patterning between deletion and normative scaling effects. Future longitudinal studies should examine how PFN allometry emerges in early life, as well as how PFN development in 22q11DS relates to psychosis-spectrum outcomes.

## Methods

### Initial Samples

Normative data were obtained from the Philadelphia Neurodevelopmental Cohort (PNC), a large community-based study of brain development (initial n=1,601)^23^. 22q11DS data were obtained from individuals with genotype-confirmed deletions evaluated by the 22q and You Center at the Children’s Hospital of Philadelphia (initial n=130)^28^.

### Image acquisition and processing

All PNC and 22q11DS MRI scans were acquired on the same 3T scanner with the same acquisition parameters^23,28^. Scans included a magnetization-prepared, rapid acquisition gradient-echo T1-weighted (MPRAGE) image as well as both resting-state (6:18 minutes) and emotional-identification task (10:36 minutes) fMRI scans. BIDS datasets were curated and validated using CuBIDS^79^, which confirmed consistent acquisition parameters across subjects. All fMRI scans were processed using fMRIPrep 25.0.0_80_ and XCP-D 0.10.6^81^.

The fMRIPrep anatomical workflow performed skull-stripping, tissue segmentation, nonlinear normalization to MNI152NLin6Asym space, and T1w-only FreeSurfer (v7.3.2) surface reconstruction^82,83^; default parameters were used throughout. The fMRIPrep functional workflow performed head motion estimation, slice-timing correction, field map-based susceptibility distortion correction, and EPI-to-T1w registration via boundary-based registration^84^, with data resampled to fsLR-32k surface space^85^. XCP-D was run in *linc* mode, which applied 36P confound regression (six motion parameters, white matter, cerebrospinal fluid, and global signal, with derivatives and quadratic terms)^86,87^, despiking, Butterworth band-pass filtering (0.01-0.08 Hz) with confounds orthogonalized to the temporal filter prior to regression^88^, and 6mm FWHM smoothing; no volume censoring was applied.

Structural image exclusions were determined by manual evaluation of T1-weighted images by expert raters^89^. Subjects with incomplete duration of either fMRI scan (less than 3 minutes) were excluded. Subjects with a median framewise displacement (FD) greater than 0.3 across both fMRI scans were also excluded to minimize motion-related confounds^90^. Filtering resulted in a normative PNC dataset of n=1,332, ages 8-23, and a 22q11DS dataset of n=100, ages 10-47. Euler number, a measure of surface reconstruction quality based on the number of topological defects^91^, was derived using freesurfer-post 0.1.2^92^ and included as a model covariate rather than an exclusion criterion.

### Demographic matching of PNC and 22q11DS individuals

To find matched controls for 22q11DS cases, we matched on the biological variables of age and sex, the societal variable of race to account for ascertainment disparities, and the technical variable of median FD to minimize motion-related confounds. Matching used nearest-neighbor matching in three serial rounds (MatchIt)^93^ on propensity scores from a logistic regression of 22q11DS status on these four variables, with a caliper of 0.1 standard deviations of the propensity score logit. Of 100 eligible cases, 62 were matched to 153 PNC controls (matched sample ages 10-24; covariate balance reported in **Supplementary Table 5**); remaining cases were primarily unmatched due to older age. The full PNC sample (n=1,332) was used for normative scaling analyses, and the matched case-control sample for 22q11DS analyses.

### Personalized Functional Network (PFN) Derivation

In both datasets, 17 PFNs were derived from concatenated rest and task fMRI data (maximum 16:54 minutes) using spatially-regularized non-negative matrix factorization (SR-NMF)^31^ via the pNet toolbox^94^. SR-NMF positively weights covarying connectivity patterns within each subject’s concatenated time series to identify individual-specific definitions of *k* = 17 networks. A group consensus atlas of networks (**Fig. 1C**; **Supplementary Fig. 1**) from past work in the PNC^7^ was used as a prior in time series decomposition, allowing for correspondence with PFN results from our previous studies^7,9,22,71^. To aid in interpretation of functional neuroanatomy, we used the Network Correspondence Toolbox^33^ to test alignment of our group atlas with four previously characterized functional network atlases (Yeo 17^95^, Gordon 17^6^, Laumann 12^5^, and Du 15^96^) and based on resulting Dice coefficients and significance testing, assigned functional names to each network (**Supplementary Table 6**).

The SR-NMF method yields a loading matrix of 17 networks by 59,412 vertices (non-medial wall vertices in fsLR 32k template), in which each value quantifies the extent to which a vertex belongs to a network, a soft parcellation of network identity. Subjects in the normative PNC sample (n=1,332) and the matched 22q11DS cases (n=62) passed pNet quality control criteria of functional coherence and spatial correspondence (**Supplementary Fig. 10**). Loading values were normalized such that, for each vertex, the loadings across the 17 networks sum to 1, giving a probabilistic network definition that we refer to as PFN topography.

### Area Calculations: Total Cortical Surface and Network Areas

Vertex-wise surface area values were calculated from each subject’s midthickness surface (resampled to fsLR 32k template space) using the Connectome Workbench command *wb_command --surface-vertex-areas*^97^. The midthickness surface was chosen as it most closely approximates the cortical depth of the fMRI signal used for functional network derivation (described above). Total cortical surface area (TSA) of each subject was then calculated by summing the vertex-wise surface area values across the cortical surface, excluding the medial wall (59,412 vertices).

The surface areas of cortical networks were calculated in two ways (**Fig. 1F**): anatomical network area, based on group-atlas-defined networks, and functional network area, based on each individual’s PFN topography. For anatomical network areas, each subject’s vertex-wise surface area values were weighted by a given network’s loadings based on the group atlas soft parcellation (**Supplementary Fig. 1**), with the resulting network-weighted surface areas summed across the cortical surface to yield a network anatomical area value (17 network values per subject). For functional network areas, a subject’s vertex-wise surface area values were weighted by network loadings from that subject’s PFN topography rather than the group atlas, and subsequently summed in the same way.

Anatomical area therefore varies across participants only through vertex-wise surface area, whereas functional area also varies through PFN topography. To further isolate topography, we computed functional extent as each participant’s PFN loadings weighted by fixed fsLR template vertex areas, indexing differences in network allocation independent of surface area. Thus, anatomical area reflects surface area variability alone, functional extent reflects topographical variability alone, and functional area reflects both.

### Statistical Analysis

Because our samples span ages 8-24, we modeled age flexibly through a generalized additive modeling approach to reflect nonlinear developmental effects. Age was included as a penalized thin plate regression spline (k=4)^98^ in generalized additive models fit with the mgcv package: network-level models used the *gam()* function^99^ with restricted maximum likelihood (REML) estimation^100^, and vertex-level models used the *bam()* function with fast REML (fREML)^101^.

To understand if functional areal scaling differs from anatomical areal scaling in typical development, the relationships of both functional and anatomical networks areas with total cortical surface area (TSA) were modeled allometrically^3,4,14^ (log-log transformed, natural log *e*) in PNC (**Model 1**). In addition to the age spline, model covariates included sex, median FD, and Euler number. The β coefficient of the log(TSA) term reflects allometry, with β=1 indicating isometry, β>1 indicating hyperallometry, and β<1 indicating hypoallometry. Wald tests were used to determine significant deviation from isometry; multiple comparisons were accounted for via Bonferroni correction across the 17 networks within each model type. Additionally, we computed the absolute difference in allometric coefficients (|β_PFN_ − β_Group_|) for each network and reported its mean and standard deviation across association and sensorimotor networks (classification in **Fig. 1C**). Because both total cortical surface area^2^ and PFN topography^7^ vary across our age range of 8-23, we assessed whether our estimates were influenced by age-related covariation by refitting models without the age spline (**Model 1a**) and comparing the resulting allometric coefficients to Model 1.

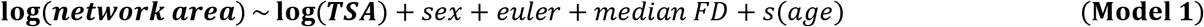

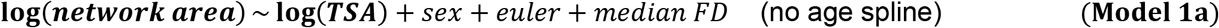

Scaling of functional extent was modeled identically as in **Model 1**. Because template vertex areas do not vary with TSA, extent coefficients index topographical scaling rather than areal allometry, with significance tested against 0 rather than 1.

Vertex-level models (**Model 2**) used the same predictors, fit to each PFN loading. Because loadings index network membership rather than area, coefficients are centered on zero rather than one. False-discovery rate (FDR) correction^102^ was applied across all variable vertex loadings. As linear-log beta coefficients can be less intuitive for interpretation, effect sizes were multiplied by ln(1.5) for visualization on cortical maps, such that transformed effects represent loading change associated with 50% increase in TSA. Vertex-level anatomical allometry models were also tested (**Model 3**), replicating prior work^3^.

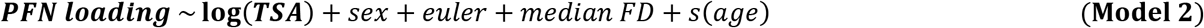

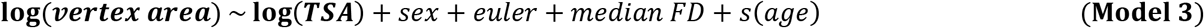

To understand if 22q11DS impacts network areas, the relationships of both anatomical and functional network areas with 22q11DS status were tested in our matched sample. We included models with and without scaling adjustment (log-transformed TSA covariate) (**Model 4**). We also included a random effect to account for case-control matching structure, utilizing *s(subclass, bs = “re”)* within *mgcv*. Other covariates, as well as multiple comparisons correction approach, were identical to the normative scaling Model 1. As network areas remained log-transformed, effect sizes were exponentiated relative to 1 (i.e., (*e*^β^-1)×100) to allow for percent change interpretation of 22q11DS effects. We also included sensitivity analyses of age covariate exclusion (**Model 4a**). Furthermore, we investigated scaling-adjusted developmental trajectories of 22q11DS PFN effects using age-by-genotype interaction models (**Model 4b**), as well as whether network scaling itself changed in 22q11DS, using scaling-by-genotype interaction models (**Model 4c**).

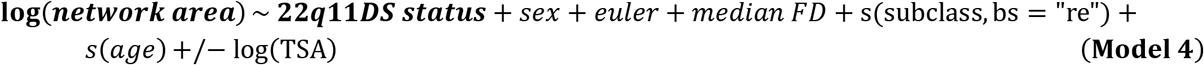

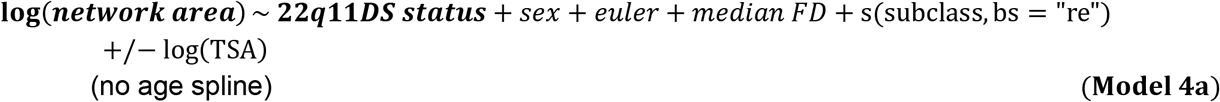

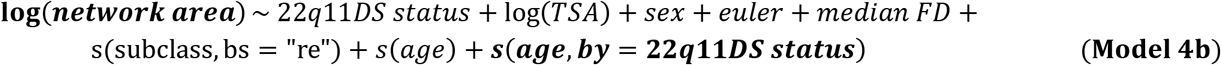

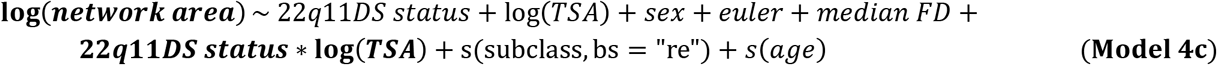

22q11DS effect on functional extent was modeled identically as in **Model 4**.

Complementary vertex-level PFN topography models were used to further interrogate 22q11DS PFN effects (**Model 5**), using identical predictors as Model 4 (with and without scaling adjustment), and then subsequently FDR-corrected across all variable vertex loadings. Vertex-level anatomical models of 22q11DS effects were also tested (**Model 6**), with effects exponentiated relative to 1 ((*e*^β^-1)×100).

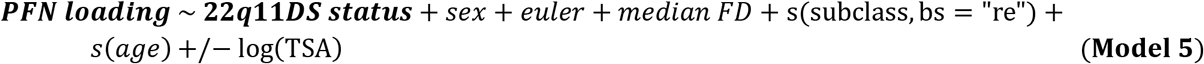

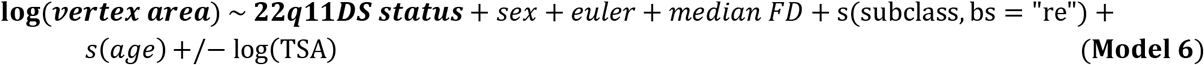

In both scaling and 22q11DS models, we tested whether functional network area effects differed from anatomical network area effects using paired bootstrap resampling (5,000 iterations). For normative scaling, subjects were resampled with replacement, and for 22q11DS, case-control subclasses were resampled with replacement to preserve matching structure. Within each bootstrap iteration, PFN and group-atlas models were refitted using the same resampled observations, and the difference between their regression coefficients was calculated. For normative scaling, we additionally calculated the difference in absolute deviation from isometry (|β_PFN_ − 1| − |β_Group_ − 1|). Uncorrected 95% bias-corrected and accelerated (BCa) bootstrap confidence intervals^103^ were estimated for individual coefficients and comparison statistics. Two-sided p-values for the comparisons were calculated by inverting the corresponding BCa confidence intervals using the *boot.pval* package, with the null hypothesis of zero difference. P-values were Bonferroni-corrected across 17 networks separately for each comparison statistic.

To assess the association between normative PFN scaling and scaling-adjusted 22q11DS differences in functional network extent (**Fig. 6B**), we also applied matched-subclass bootstrap resampling (5,000 iterations), refitting 22q11DS-effect models for all 17 networks using the same resampled observations within each iteration. Pearson correlations between the resulting 22q11DS coefficients and fixed normative scaling coefficients were recalculated for each iteration, with 95% BCa confidence intervals and two-sided p-values estimated as described above, testing against a null hypothesis of zero correlation.

To visualize shifts in PFN topography across brain size variation, model-based predictions were generated for each PFN vertex-level model at 100 evenly-spaced values of log-transformed TSA spanning the 1^st^ to 99^th^ percentiles of PNC sample distribution, holding all covariates constant at average reference values. For each of the 100 TSA reference values, predicted loadings were then used to generate a hard-parcellated map using a winner-take-all approach.

Whole cortex effect size maps of PFN topography (**Figs. 3C, 6A**) display the maximum magnitude across all 17 network effect sizes for each cortical vertex. PFN topography variability was derived using the median absolute deviation (MAD) of each PFN loading value across individuals in the PNC cohort (n=1,332) and averaging those MAD values across the 17 networks for each cortical vertex. Spatial correspondence across cortical maps was tested using the *neuromaps* toolbox^104^ and spin test approach (N=5000)^105^. Map visualizations of vertex-level scaling and 22q11DS results were rendered via Connectome Workbench^97^.

