## Supplementary Data for "Personalized functional networks scale with brain size beyond anatomy and are reorganized in 22q11.2 deletion syndrome"

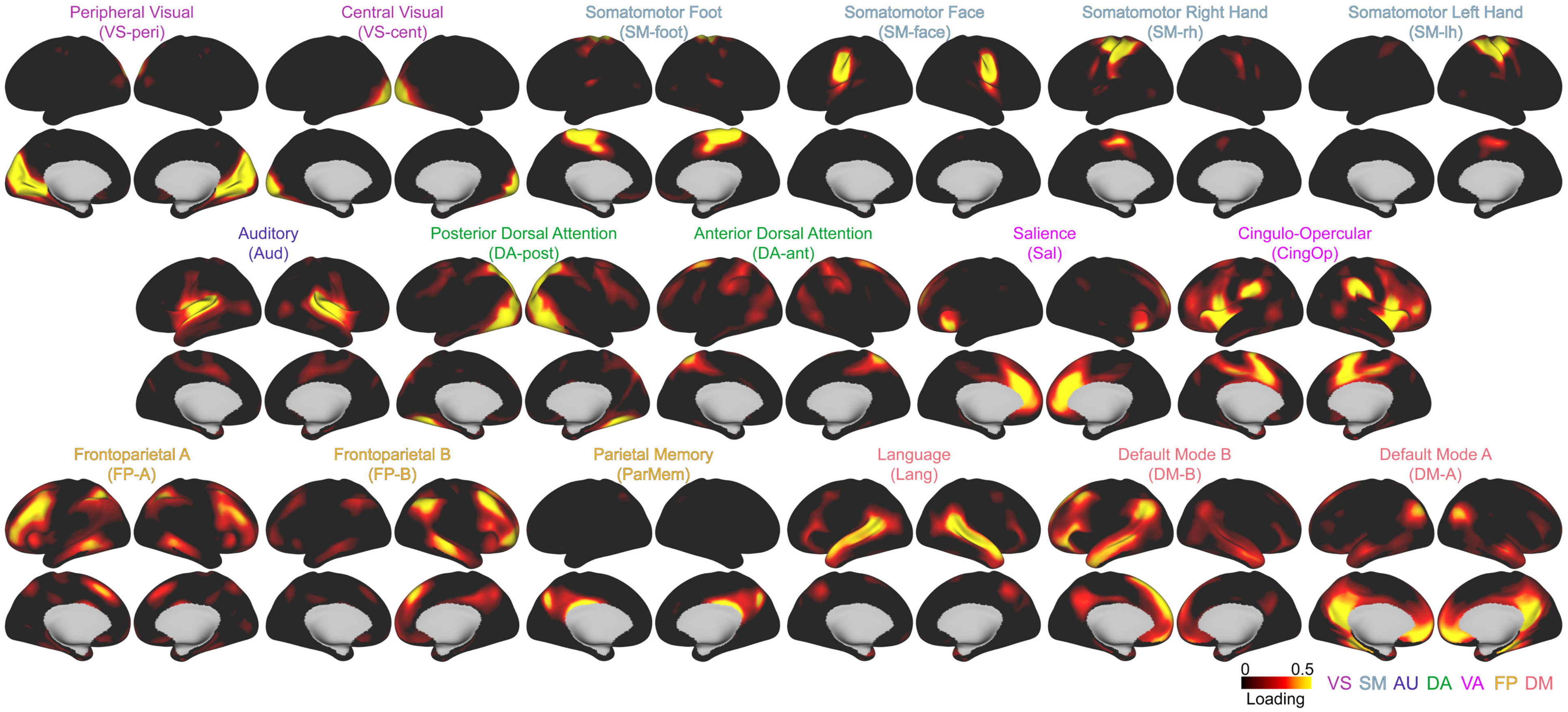


***Supplementary Figure 1. Group atlas network loading topography maps.*** Network maps of group atlas loading topography, as defined in prior work[1,2]. These maps were used in the calculation of anatomical network area (Fig. 1F). Network labels determined in Supplementary Table 6, system-level categories used for color-coding here. **Functional systems**: VS = visual; SM = somatomotor; AU = auditory; DA = dorsal attention; VA = ventral attention; FP = frontoparietal; DM = default mode.

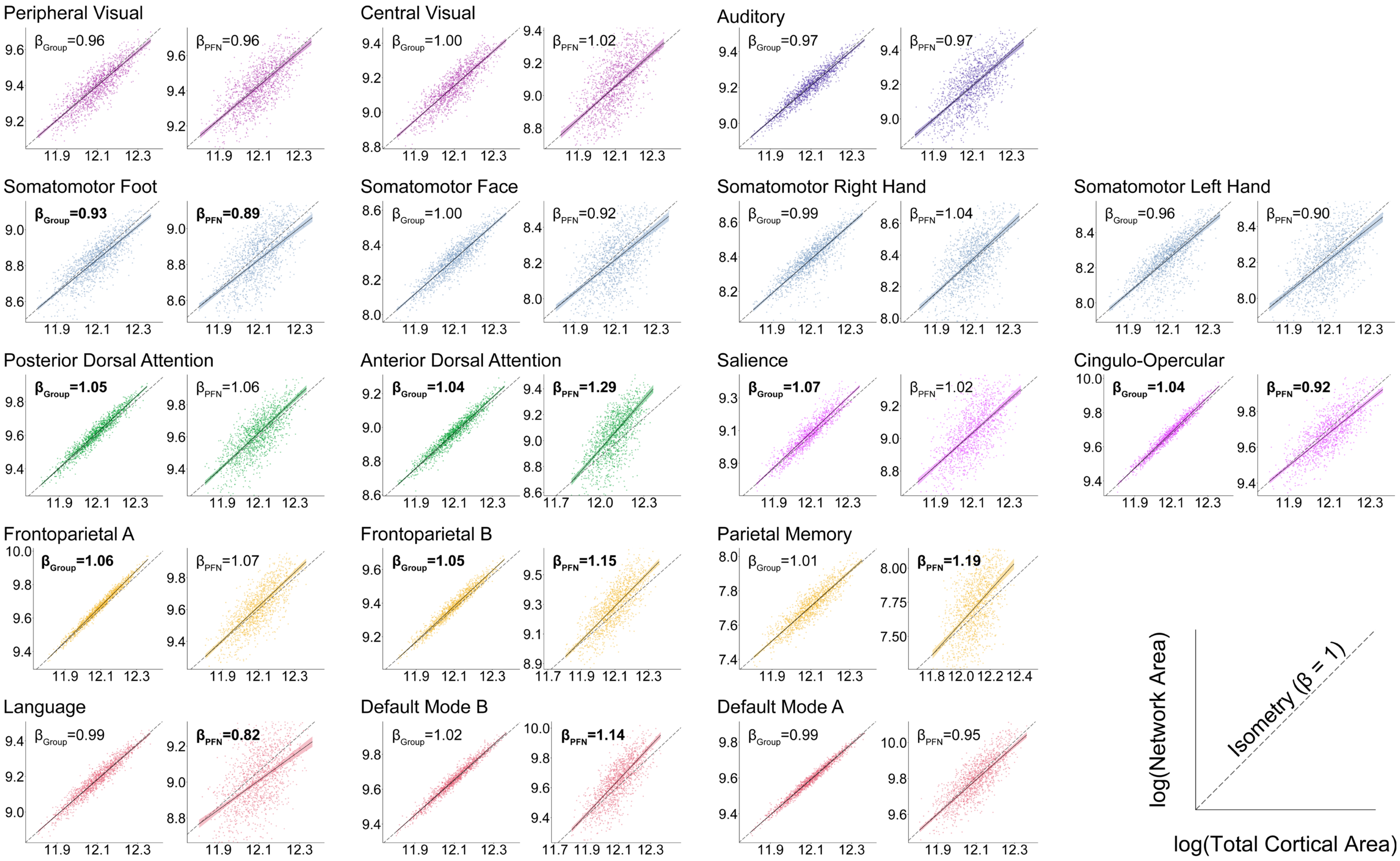


***Supplementary Figure 2. Log-log plots of group-atlas and personalized network area allometry.*** Log-log plots show anatomical group-atlas (left side) and functional PFN-based network (right side) areal allometry based on individual-level observations (points) and model-predicted scaling trajectories across the observed range of total cortical area (lines), with covariates held at representative values based on marginal distributions. Dotted line included as an isometry reference, anchored to the model-predicted network area at the minimum observed total cortical area. Bolded beta annotations indicate significant deviation from isometry after multiple comparisons correction within model type across the 17 networks ($p_{Bonf}$ < 0.05).

| | Group Atlas | | PFNs | | Functional Deviation (PFNs vs Group Atlas) | | | | Functional Extent | |
|---|---|---|---|---|---|---|---|---|---|---|
| | β | $p_{Bonf}$ | β | $p_{Bonf}$ | $\beta_{PFN} - \beta_{Group}$ | $p_{Bonf}$ | $\lvert\beta_{PFN} - 1\rvert - \lvert\beta_{Group} - 1\rvert$ | $p_{Bonf}$ | β | $p_{Bonf}$ |
| VS-peri | 0.96 | 0.88 | 0.96 | 1 | −0.01 | 1 | 0.01 | 1 | 0.00 | 1 |
| VS-cent | 1.00 | 1 | 1.02 | 1 | 0.02 | 1 | 0.02 | 1 | 0.02 | 1 |
| SM-foot | 0.93 | 0.01 | 0.89 | 0.03 | −0.04 | 1 | 0.04 | 1 | −0.04 | 1 |
| SM-face | 1.00 | 1 | 0.92 | 0.71 | −0.08 | 0.55 | 0.08 | 0.47 | −0.08 | 0.44 |
| SM-rh | 0.99 | 1 | 1.04 | 1 | 0.05 | 1 | 0.04 | 1 | 0.05 | 1 |
| SM-lh | 0.96 | 1 | 0.90 | 0.37 | −0.06 | 1 | 0.06 | 1 | −0.05 | 1 |
| Auditory | 0.97 | 0.09 | 0.97 | 1 | 0 | 1 | 0.00 | 1 | 0.01 | 1 |
| DA-post | 1.05 | $2.70\times10^{-6}$ | 1.06 | 0.6 | 0.01 | 1 | 0.01 | 1 | 0.00 | 1 |
| DA-ant | 1.04 | $1.20\times10^{-4}$ | 1.29 | $4.50\times10^{-9}$ | 0.24 | $3.40\times10^{-3}$ | 0.24 | $3.40\times10^{-3}$ | 0.24 | $2.10\times10^{-6}$ |
| Salience | 1.07 | $3.00\times10^{-6}$ | 1.02 | 1 | −0.05 | 1 | −0.05 | 0.37 | −0.06 | 1 |
| CingOp | 1.04 | $2.20\times10^{-6}$ | 0.92 | 0.01 | −0.12 | $3.40\times10^{-3}$ | 0.05 | 1 | −0.12 | $3.40\times10^{-6}$ |
| FP-A | 1.06 | $<1.0\times10^{-16}$ | 1.07 | 0.35 | 0 | 1 | 0.00 | 1 | 0.00 | 1 |
| FP-B | 1.05 | $7.60\times10^{-11}$ | 1.15 | $2.60\times10^{-5}$ | 0.1 | 0.03 | 0.10 | 0.03 | 0.09 | 0.06 |
| ParMem | 1.01 | 1 | 1.19 | 0.02 | 0.18 | 0.04 | 0.18 | 0.04 | 0.19 | 0.01 |
| Language | 0.99 | 1 | 0.82 | $6.00\times10^{-4}$ | −0.17 | $3.40\times10^{-3}$ | 0.17 | $3.40\times10^{-3}$ | −0.16 | $2.00\times10^{-3}$ |
| DM-B | 1.02 | 0.26 | 1.14 | $7.10\times10^{-5}$ | 0.12 | 0.01 | 0.12 | 0.01 | 0.11 | $1.40\times10^{-3}$ |
| DM-A | 0.99 | 0.25 | 0.95 | 0.39 | −0.04 | 1 | 0.04 | 1 | −0.04 | 1 |

***Supplementary Table 1. Table of all network-level scaling effects in the Philadelphia Neurodevelopmental Cohort.*** Effect sizes and multiple comparisons-corrected significance values of group atlas areal allometry, PFN areal allometry, difference between the two, difference in deviation from isometry between the two, and functional extent scaling. Color of effect size cells indicates relative degree of hyperallometry ($\beta > 1$; red) and hypoallometry ($\beta < 1$; blue) across group atlas and PFN allometry; functional extent color range is separate, with values centered around 0 instead of 1. Functional extent effects capture scaling attributable to network topography alone independent of vertex-level surface area variation, and thus functional extent effects were nearly identical to functional deviation in scaling ($\beta_{PFN}$ - $\beta_{Group}$). **Network abbreviations**: CingOp = cingulo-opercular; DA-ant = anterior dorsal attention; DA-post = posterior dorsal attention; ParMem = parietal memory; SM-lh = left hand; SM-rh = right hand; VS-cent = central visual; VS-peri = peripheral visual.

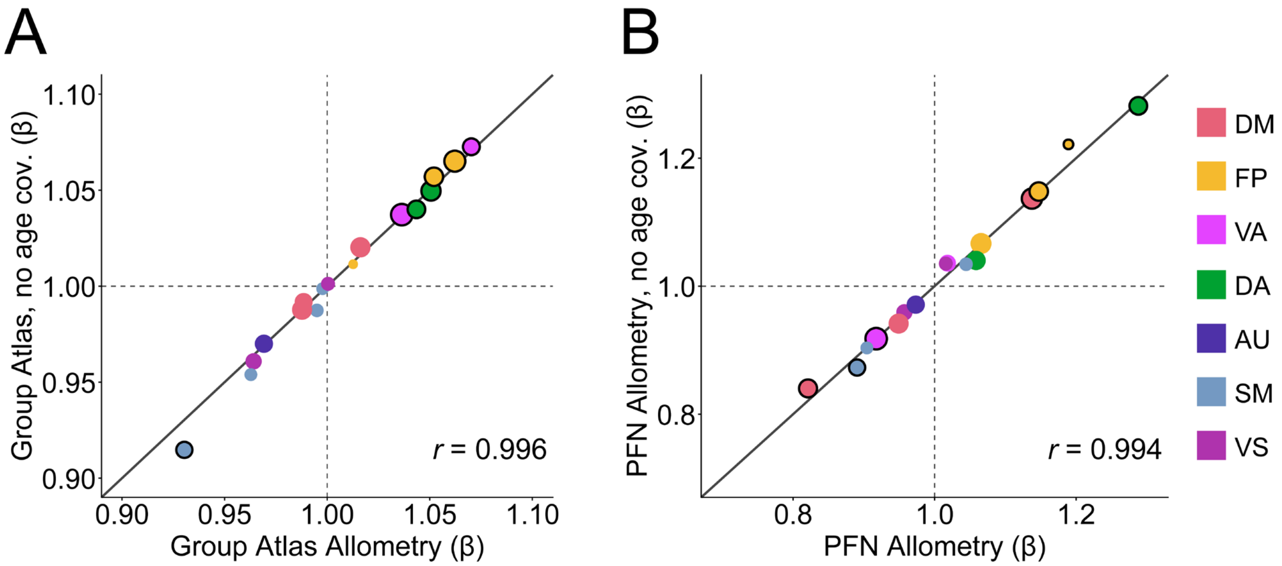


***Supplementary Figure 3. Network area allometry is not dependent on age covariate specification.*** Allometric beta coefficients in models with and without age spline covariate inclusion show highly consistent results in both group-atlas-defined networks (**A**) and PFNs (**B**). Diagonal line represents equality between the effects of the two different model specifications. Black outlines indicate significant allometry coefficients, which were found to be identical with and without age spline inclusion. Size of points reflect group atlas network size. **Functional systems**: DM = default mode; FP = frontoparietal; VA = ventral attention; DA = dorsal attention; AU = auditory; SM = somatomotor; VS = visual.

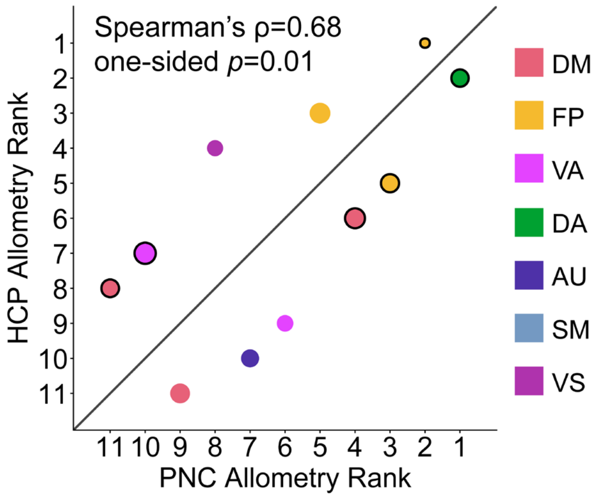


| Cui-17 Networks | Yeo-17 Match | Dice | $p_{Bonf}$ | PNC allometry rank | HCP allometry rank |
|---|---|---|---|---|---|
| VS-peri | Visual B | 0.92 | 0.02 | 9 | 4 |
| VS-cent | Visual A | 0.67 | 0.07 | - | - |
| SM-foot | SM A | 0.68 | 0.07 | - | - |
| SM-face | SM B | 0.48 | 0.14 | - | - |
| SM-rh | SM A | 0.31 | 0.44 | - | - |
| SM-lh | SM A | 0.33 | 0.46 | - | - |
| Auditory | SM B | 0.72 | 0.02 | 8 | 10 |
| ***DA-post*** | ***Dorsal A*** | ***0.70*** | ***0.02*** | 6 | 12 |
| DA-ant | Dorsal B | 0.41 | 0.02 | 1 | 2 |
| Salience | Sal/Ventral B | 0.42 | 0.03 | 7 | 9 |
| CingOp | Sal/Ventral A | 0.67 | 0.02 | 11 | 7 |
| FP-A | Control A | 0.65 | 0.02 | 5 | 3 |
| FP-B | Control B | 0.54 | 0.02 | 3 | 5 |
| ParMem | Control C | 0.48 | 0.03 | 2 | 1 |
| Language | Temp Par | 0.69 | 0.03 | 12 | 8 |
| DM-B | Default B | 0.63 | 0.03 | 4 | 6 |
| DM-A | Default A | 0.50 | 0.02 | 10 | 11 |

| Yeo-17 Networks | Kong-17 Match | Dice | $p_{Bonf}$ |
|---|---|---|---|
| Control A | Control A | 0.49 | 0.02 |
| Control B | Control B | 0.52 | 0.02 |
| Control C | Control C | 0.33 | 0.02 |
| Default A | Default A | 0.47 | 0.03 |
| Default B | Default B | 0.61 | 0.02 |
| Default C | Default C | 0.53 | 0.02 |
| Dorsal B | Dorsal B | 0.56 | 0.02 |
| ***Dorsal A*** | ***Visual A*** | ***0.44*** | ***0.02*** |
| Limbic A | Default A | 0.26 | 0.14 |
| Limbic B | Control B | 0.20 | 0.39 |
| Sal/Ventral A | Sal/Ventral A | 0.65 | 0.02 |
| Sal/Ventral B | Sal/Ventral B | 0.40 | 0.03 |
| SM A | SM A | 0.78 | 0.02 |
| SM B | SM B | 0.60 | 0.03 |
| Temp Par | Language | 0.32 | 0.02 |
| Visual A | Visual C | 0.44 | 0.03 |
| Visual B | Visual B | 0.81 | 0.02 |

***Supplementary Figure 4. Correlation of homologous networks reveals consistent PFN areal allometry between independent studies.*** Plot shows correlation of effect rank between our findings with a prior report of functional network allometry in the Human Connectome Project (HCP)[3]. A significant positive correlation was found in PFN allometry relative ordering defined in development (PNC) versus adulthood (HCP): Spearman's ρ=0.68, one-sided *p*=0.01. Diagonal line represents equality between study effect ranks; black outlines indicate significant allometry coefficients in our results. To facilitate cross-study comparison of results, we identified homologous networks using the network correspondence toolbox. The HCP study used vertex-level Kong-17 networks[4] as their group atlas prior. Because this vertex-level atlas was not available in the toolbox, network identities were harmonized using the Yeo-17 atlas as a common reference[5]. To establish homologous networks between our group atlas (Cui-17) and the Yeo-17 atlas, we retained the highest Dice correspondence among significant matches ($p_{Bonf} < 0.05$), shown in the middle table. Correspondence between Yeo-17 and Schaefer-based parcel-level[6] Kong-17 networks were also evaluated to facilitate interpretation of the published findings, showing strong consistency across the vast majority of networks as displayed in the right table. Analyses were restricted to networks exhibiting unique, significant network matches across both comparisons (non-significant matches grayed out). The posterior dorsal attention network (DA-post) was excluded from the primary analysis as its Yeo-17 counterpart (Dorsal Attention A) mapped most strongly to Kong-17 Visual A, representing a discrepancy in broad functional-system classification (highlighted cells). This procedure yielded 11 matched networks. As a sensitivity analysis, we included the posterior dorsal attention network, using the allometry rank of the Kong-17 homologous network for HCP allometry (Visual A network), resulting in a consistent correlation: Spearman's ρ=0.59, one-sided *p*=0.02. **Network abbreviations**: CingOp = cingulo-opercular; DA-ant = anterior dorsal attention; DA-post = posterior dorsal attention; ParMem = parietal memory; Sal = salience; SM-lh = left hand; SM-rh = right hand; Temp Par = temporal parietal; VS-cent = central visual; VS-peri = peripheral visual.

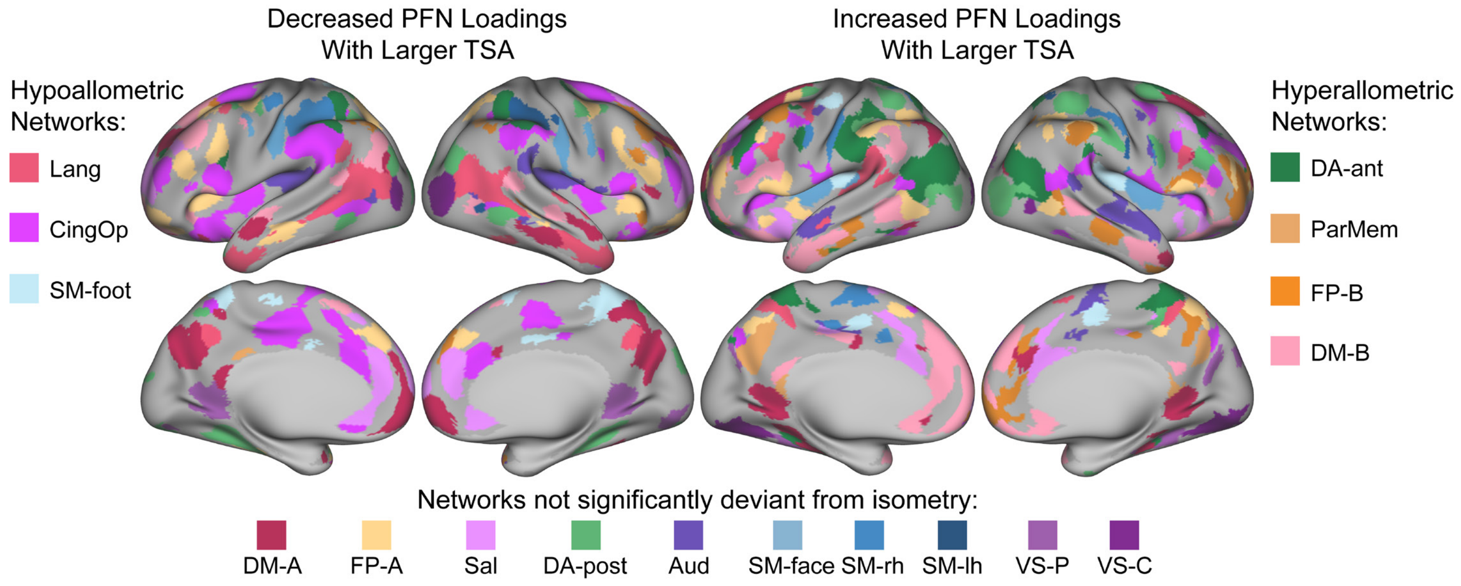


***Supplementary Figure 5. Directional vertex-level PFN loading associations with cortical scaling in typical development.*** Significant ($p_{FDR} < 0.05$) negative scaling effects (i.e., associated with cortical territory loss) and positive scaling effects (i.e., associated with cortical territory gain) of all networks (extension of Fig. 3A). 50 $mm^2$ cluster threshold applied; in cases of overlapping network vertices, the network with the larger absolute effect size is displayed. Hypo- and hyperallometric network categorization reflects PFN areal allometry as shown in Fig. 2B. **Network abbreviations**: Aud = auditory; CingOp = cingulo-opercular; DA-ant = anterior dorsal attention; DA-post = posterior dorsal attention; DM = default mode; FP = frontoparietal; Lang = language; ParMem = parietal memory; Sal = salience; SM = somatomotor; SM-lh = left hand; SM-rh = right hand; VS-C = central visual; VS-P = peripheral visual.

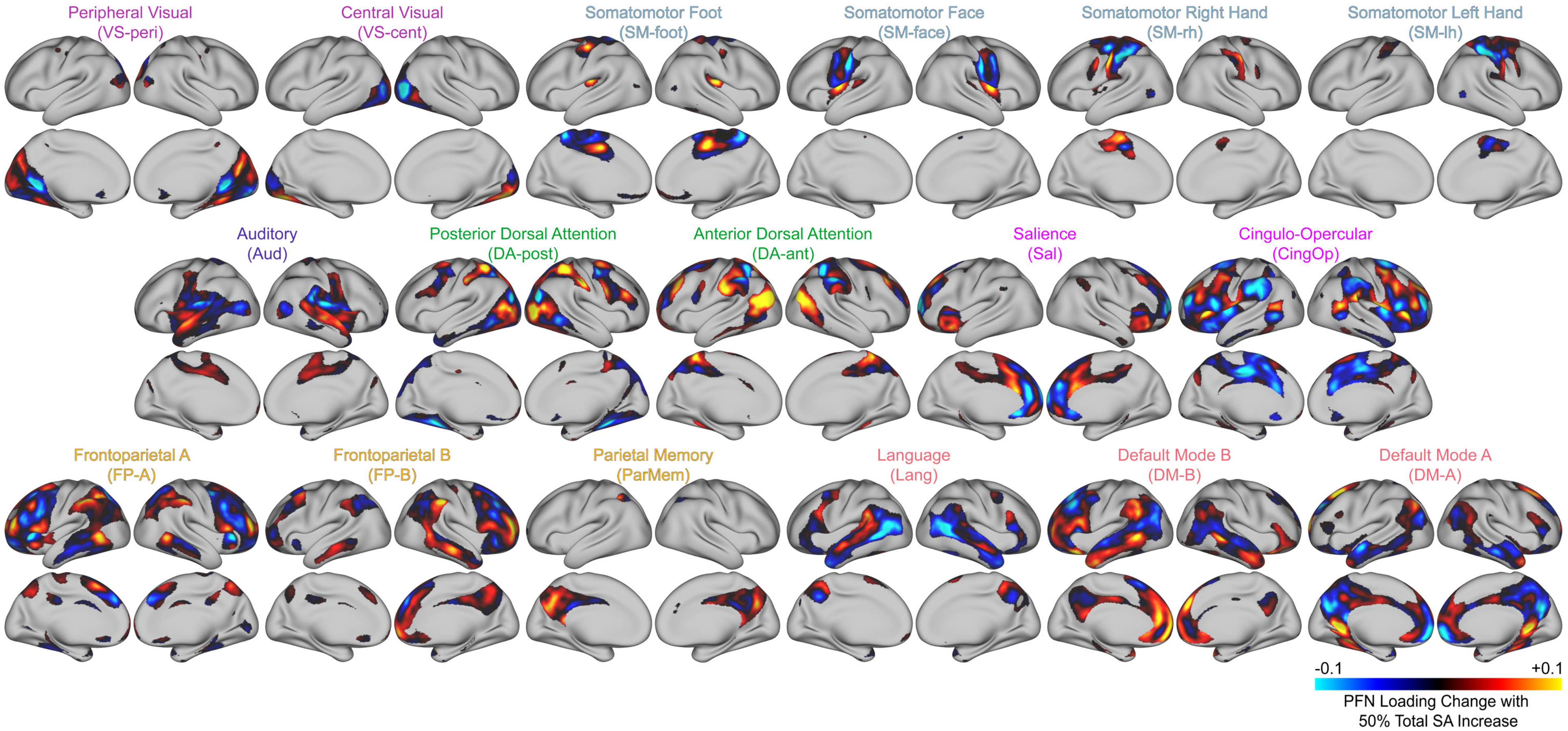


***Supplementary Figure 6. Unthresholded vertex-level PFN loading changes with cortical scaling in typical development.*** 17 network effect size maps of PFN loading change with increase in total surface area (SA), visualized as the change in PFN loading associated with a 50% increase in total cortical surface area. The absolute maximum value across these 17 networks is displayed in Fig. 3C.

| | Scaling Unadjusted | | | | | | Scaling Adjusted | | | | | | | |
|---|---|---|---|---|---|---|---|---|---|---|---|---|---|---|
| | Group Atlas | | PFN | | Functional Deviation (PFN vs Group Atlas) | | Group Atlas | | PFN | | Functional Deviation (PFN vs Group Atlas) | | Functional Extent | |
| | % Change | $p_{Bonf}$ | % Change | $p_{Bonf}$ | PFN - Group | $p_{Bonf}$ | % Change | $p_{Bonf}$ | % Change | $p_{Bonf}$ | PFN - Group | $p_{Bonf}$ | % Change | $p_{Bonf}$ |
| VS-peri | −9.6% | $1.52\times10^{-9}$ | −10.3% | $8.15\times10^{-8}$ | −0.7% | 1 | -5.2% | $2.72\times10^{-7}$ | -5.9% | $1.52\times10^{-4}$ | −0.8% | 1 | −0.7% | 1 |
| VS-cent | −9.1% | $1.12\times10^{-8}$ | −7.5% | 0.03 | 1.7% | 1 | -4.7% | $2.58\times10^{-6}$ | −2.3% | 1 | 2.4% | 1 | 2.5% | 1 |
| SM-foot | -5.1% | $4.72\times10^{-3}$ | −3.6% | 0.96 | 1.7% | 1 | −0.5% | 1 | 1.4% | 1 | 1.9% | 1 | 0.7% | 1 |
| SM-face | −1.6% | 1 | −1.2% | 1 | 0.4% | 1 | 2.9% | $2.8\times10^{-3}$ | 2.8% | 1 | −0.1% | 1 | 0.1% | 1 |
| SM-rh | -4.7% | 0.03 | -4.9% | 0.54 | −0.2% | 1 | 0.1% | 1 | 0.5% | 1 | 0.5% | 1 | 0.9% | 1 |
| SM-lh | -4.1% | 0.24 | −0.4% | 1 | 3.8% | 1 | 0.3% | 1 | 4.3% | 0.72 | 4.0% | 1 | 4.6% | 0.41 |
| Auditory | −2.1% | 1 | -4.5% | 0.30 | −2.5% | 1 | 2.6% | $1.33\times10^{-4}$ | 0.2% | 1 | −2.4% | 1 | −2.6% | 1 |
| DA-post | −7.4% | $1.24\times10^{-6}$ | −7.3% | $8.25\times10^{-4}$ | 0.1% | 1 | −2.4% | $1.11\times10^{-5}$ | −2.2% | 1 | 0.2% | 1 | 0.2% | 1 |
| DA-ant | -4.9% | $4.76\times10^{-3}$ | −10.8% | $1.92\times10^{-4}$ | −6.1% | 0.05 | 0.2% | 1 | -4.5% | 0.38 | -4.8% | 0.31 | -4.5% | 0.35 |
| Salience | -4.1% | 0.06 | −1.2% | 1 | 3.0% | 1 | 1.1% | 0.73 | 4.7% | 0.27 | 3.5% | 1 | 3.6% | 0.95 |
| CingOp | −3.1% | 0.24 | −0.5% | 1 | 2.7% | 0.35 | 1.8% | $7.79\times10^{-7}$ | 4.3% | $4.61\times10^{-3}$ | 2.5% | 0.56 | 2.2% | 0.77 |
| FP-A | -4.3% | 0.02 | −1.3% | 1 | 3.1% | 0.83 | 0.7% | 0.05 | 2.7% | 0.71 | 2.0% | 1 | 2.0% | 1 |
| FP-B | −3.7% | 0.06 | −6.6% | $3.76\times10^{-3}$ | −3.0% | 0.90 | 1.3% | $7.78\times10^{-4}$ | −2.3% | 1 | −3.6% | 0.39 | -4.0% | 0.07 |
| ParMem | −6.2% | $2.83\times10^{-4}$ | -9.8% | 0.03 | −3.8% | 1 | −1.2% | 1 | -3.4% | 1 | −2.3% | 1 | −1.9% | 1 |
| Language | −2.8% | 0.40 | −2.0% | 1 | 0.8% | 1 | 1.7% | 0.02 | 1.0% | 1 | −0.7% | 1 | −0.8% | 1 |
| DM-B | −3.4% | 0.10 | -4.8% | 0.22 | −1.5% | 1 | 1.4% | $9.47\times10^{-5}$ | 0.9% | 1 | −0.4% | 1 | −0.8% | 1 |
| DM-A | -4.6% | $2.27\times10^{-3}$ | -5.2% | 0.01 | −0.6% | 1 | 0.1% | 1 | −0.3% | 1 | −0.5% | 1 | −0.4% | 1 |

***Supplementary Table 2. Table of all network-level 22q11DS effects before and after scaling adjustment.*** Effect sizes and multiple comparisons-corrected significance values of group atlas area effects, PFN area effects, difference between the group atlas and PFN effects, as well as functional extent effects after scaling adjustment. As network areas were log-transformed and predictors were linear (22q11DS genotype), β were transformed ($(e^{\beta}-1)\times100$) to allow for percent change effect size interpretation. Color of effect size cells indicates relative % change across all effect sizes (red indicates expansion, blue indicates reduction). Number of significant effects were attenuated after accounting for both scaling adjustment and personalized network definitions (9 group atlas scaling unadjusted; 2 PFN scaling adjusted). **Network abbreviations**: CingOp = cingulo-opercular; DA-ant = anterior dorsal attention; DA-post = posterior dorsal attention; ParMem = parietal memory; SM-lh = left hand; SM-rh = right hand; VS-cent = central visual; VS-peri = peripheral visual.

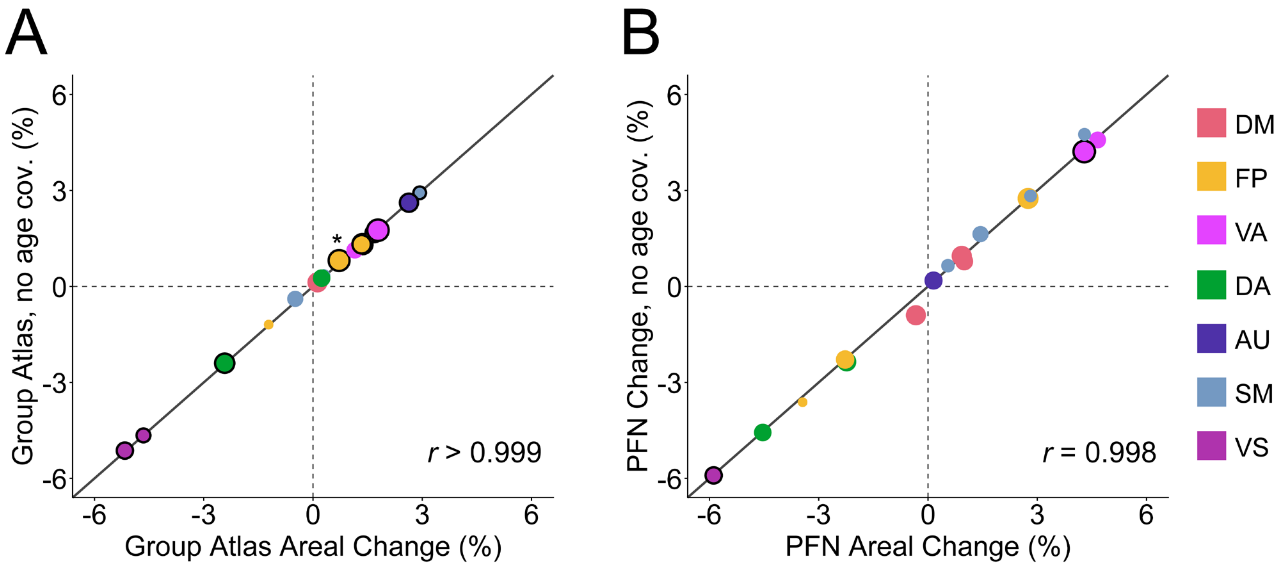


***Supplementary Figure 7. Scaling-adjusted 22q11.2 deletion effects on network areas are not dependent on age covariate specification.*** 22q11.2 deletion (22q11DS) areal effects relative to total cortical size in models with and without age spline covariate inclusion show highly consistent results in both group-atlas-defined networks (**A**) and PFNs (**B**). Diagonal line represents equality between the effects of the two different model specifications. Black outlines indicate significant 22q11DS effects in models without the age covariate, which were found to be highly consistent with and without age spline inclusion, the single exception being a relative expansion of the group-atlas-defined frontoparietal A network (asterisked) that was only found to be significant when the age spline covariate was excluded. Size of points reflect group atlas network size. **Functional systems**: DM = default mode; FP = frontoparietal; VA = ventral attention; DA = dorsal attention; AU = auditory; SM = somatomotor; VS = visual.

| | Age x Genotype | | | | Scaling x Genotype | | | |
|---|---|---|---|---|---|---|---|---|
| | Group Atlas | | PFNs | | Group Atlas | | PFNs | |
| | F-statistic | $p_{Bonf}$ | F-statistic | $p_{Bonf}$ | β | $p_{Bonf}$ | β | $p_{Bonf}$ |
| VS-peri | 2.85 | 0.83 | 1.80 | 1 | 0.10 | 1 | 0.08 | 1 |
| VS-cent | 2.44 | 1 | 0.70 | 1 | 0.11 | 1 | 0.04 | 1 |
| SM-foot | 0.30 | 1 | 0.08 | 1 | 0.04 | 1 | −0.16 | 1 |
| SM-face | 4.20 | 0.71 | 0.70 | 1 | −0.07 | 1 | −0.18 | 1 |
| SM-rh | 0.71 | 1 | 0.06 | 1 | −0.04 | 1 | −0.11 | 1 |
| SM-lh | 0.03 | 1 | 0.03 | 1 | −0.06 | 1 | 0.24 | 1 |
| Auditory | 6.81 | 0.17 | 2.19 | 1 | −0.06 | 1 | 0.37 | 0.24 |
| DA-post | 3.48 | 1 | 0.57 | 1 | 0.08 | 1 | 0.27 | 0.66 |
| DA-ant | 0.00 | 1 | 0.85 | 1 | 0.04 | 1 | 0.05 | 1 |
| Salience | 0.26 | 1 | 1.24 | 1 | −0.01 | 1 | 0.48 | 0.15 |
| CingOp | 3.55 | 0.29 | 10.04 | 0.03 | −0.07 | 0.32 | 0.04 | 1 |
| FP-A | 1.11 | 1 | 0.98 | 1 | 0.00 | 1 | −0.42 | 0.02 |
| FP-B | 1.39 | 1 | 1.05 | 1 | −0.01 | 1 | −0.08 | 1 |
| ParMem | 0.39 | 1 | 2.51 | 1 | −0.08 | 1 | −0.06 | 1 |
| Language | 4.09 | 0.76 | 0.34 | 1 | −0.01 | 1 | −0.25 | 1 |
| DM-B | 0.51 | 1 | 2.63 | 1 | −0.02 | 1 | −0.13 | 1 |
| DM-A | 0.73 | 1 | 5.05 | 0.07 | 0.01 | 1 | 0.00 | 1 |

***Supplementary Table 3. Interaction effects of 22q11DS genotype with age- and scaling- on group-atlas network and PFN areas.*** Significant interactions include the cingulo-opercular network for age-by-genotype and the frontoparietal A network for scaling-by-genotype models, multiple-comparison corrected within model type across the 17 networks ($p_{Bonf} < 0.05$; interactions visualized in **Extended Data Fig. 3**). **Network abbreviations**: CingOp = cingulo-opercular; DA-ant = anterior dorsal attention; DA-post = posterior dorsal attention; ParMem = parietal memory; SM-lh = left hand; SM-rh = right hand; VS-cent = central visual; VS-peri = peripheral visual.

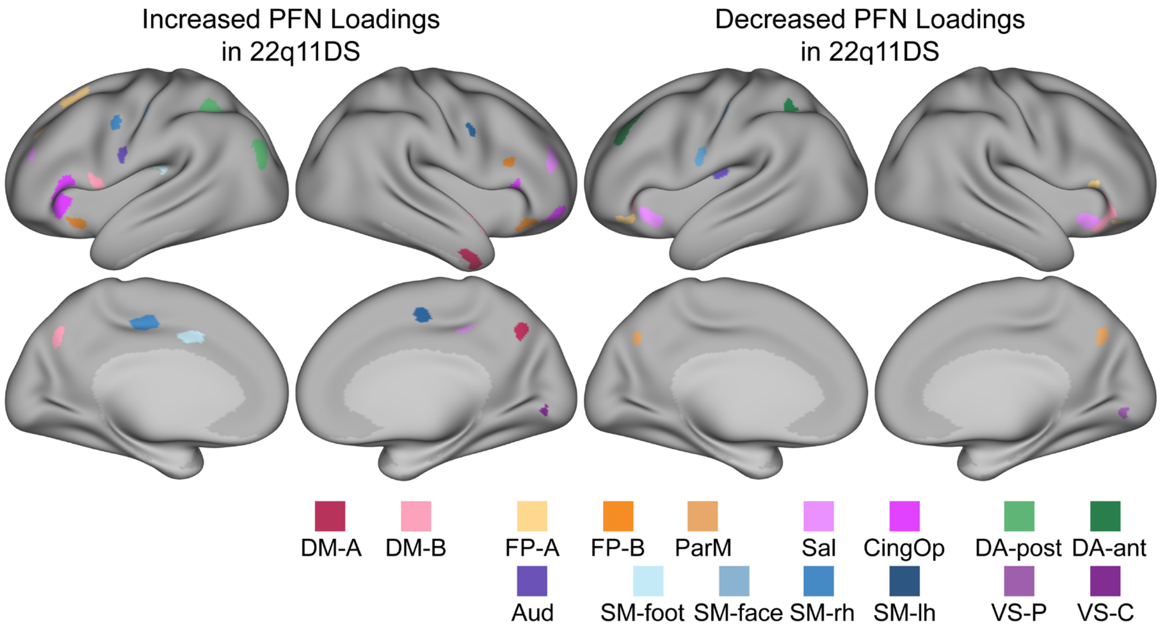


***Supplementary Figure 8. Directional vertex-level PFN loading associations with 22q11DS prior to scaling adjustment.*** Significant ($p_{FDR}$ < 0.05) positive and negative 22q11DS network loading effects prior to scaling adjustment (after scaling adjustment is shown in Fig. 5A). 50 $mm^2$ cluster threshold applied; in cases of overlapping network vertices, the network with largest effect size is displayed. Directional pattern of significant loading changes across networks remained broadly consistent before and after scaling adjustment (Fig. 5A), albeit expanded in number prior to adjustment. **Network abbreviations**: Aud = auditory; CingOp = cingulo-opercular; DA-ant = anterior dorsal attention; DA-post = posterior dorsal attention; DM = default mode; FP = frontoparietal; Lang = language; ParMem = parietal memory; Sal = salience; SM = somatomotor; SM-lh = left hand; SM-rh = right hand; VS-C = central visual; VS-P = peripheral visual.

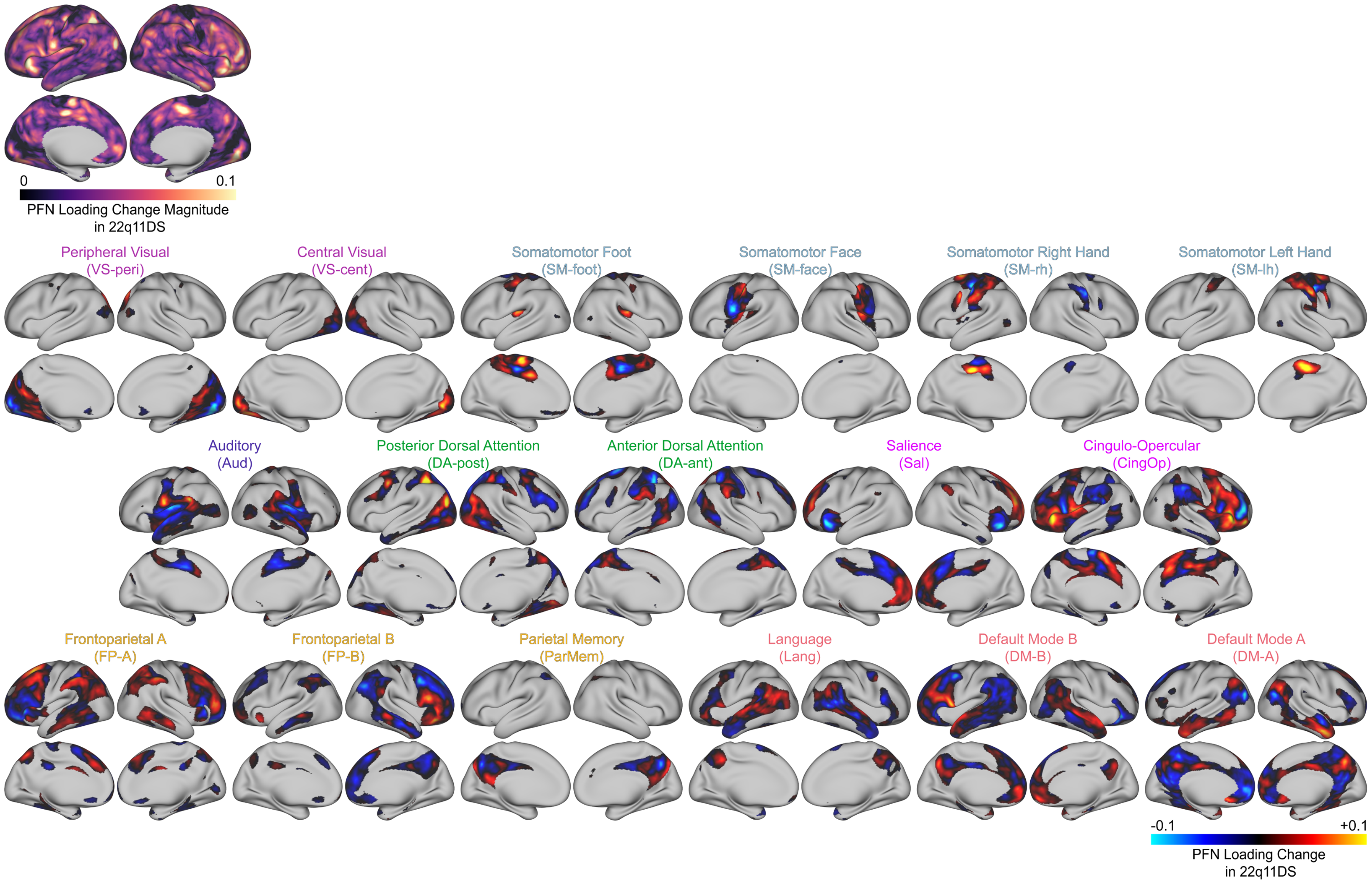


***Supplementary Figure 9. Unthresholded vertex-level PFN loading changes in 22q11DS.*** Map of the maximum absolute 22q11DS effect size across networks shown at the top, with the constituent 17 network effect size maps of PFN loading change in 22q11DS following.

| PFN vertex-level effect size maps: | Principal Gradient | | Gene Expression | | Myelination (T1/T2) | | Cortical Thickness | | Neurosynth | | Gluc. Metabolism | | Cerebral Blood Flow | | Evolutionary Exp. | |
|---|---|---|---|---|---|---|---|---|---|---|---|---|---|---|---|---|
| | *r* | $p_{Bonf}$ | *r* | $p_{Bonf}$ | *r* | $p_{Bonf}$ | *r* | $p_{Bonf}$ | *r* | $p_{Bonf}$ | *r* | $p_{Bonf}$ | *r* | $p_{Bonf}$ | *r* | $p_{Bonf}$ |
| Cortical Scaling | 0.10 | 1 | −0.17 | 0.84 | −0.27 | 0.01 | 0.05 | 1 | −0.01 | 1 | 0.00 | 1 | 0.08 | 1 | 0.01 | 1 |
| 22q11DS | 0.12 | 1 | −0.17 | 1 | −0.21 | 0.10 | 0.11 | 1 | 0.01 | 1 | 0.12 | 1 | 0.18 | 0.13 | 0.02 | 1 |

***Supplementary Table 4. Spatial correspondence of vertex-level PFN loading effect size maps with maps of cortical hierarchy.*** For biological interpretation of PFN vertex-level scaling and 22q11DS effects, we investigated the spatial correspondence of effect size maps with 8 maps of cortical hierarchy: the principal gradient of functional connectivity[7], the PC1 of gene expression[8,9], T1/T2 ratio[10], cortical thickness[10], PC1 of Neurosynth terms[11,12], glucose metabolism[13], cerebral blood flow[14], and evolutionary expansion[15]. Spatial correspondence was tested using the *neuromaps* toolbox[16] and spin test approach (N=5000)[17]. Only the map of T1/T2 ratio, a proxy for intracortical myelin, showed a significant effect with the normative scaling effect map ($r$ = −0.27, $p_{Bonf}$ = 0.01), but did not survive residualization for PFN variability ($r$ = −0.08, $p_{Bonf}$ = 1), with PFN variability defined as the inter-individual median absolute deviation averaged across the 17 network loadings in the Philadelphia Neurodevelopmental Cohort sample ($n$ = 1,332).

| | PNC controls | 22q11DS cases | *p*-value | Absolute SMD |
|---|---|---|---|---|
| Number of subjects | 153 | 62 | | |
| Age (years), mean ± SD | 18.42±3.16 | 19.12±3.69 | 0.19 | 0.20 |
| Female (%) | 45.8% | 45.2% | 0.94 | 0.01 |
| Race- White (%) | 61.4% | 66.1% | 0.52 | 0.10 |
| Race- Black (%) | 24.8% | 21.0% | 0.55 | 0.09 |
| Race- Other (%) | 13.7% | 12.9% | 0.87 | 0.02 |
| Median FD (mm), mean ± SD | 0.11±0.07 | 0.11±0.05 | 0.72 | 0.05 |

***Supplementary Table 5. Statistics on matched sample of PNC and 22q11.2 deletion syndrome participants.*** PNC controls were selected using an up-to-3:1 matching scheme on age, sex, race, and median framewise displacement (FD). Case-control models using this sample included sex, median FD, and Euler number as linear covariates, matching subclass as a random effect, and an age spline. SMD = standardized mean difference; FD = framewise displacement.

| Cui-17 Original | | Yeo-17 | | | Gordon-17 | | | Du-15 | | | Laumann-12 | | | Cui-17 Renamed Network Labels |
|---|---|---|---|---|---|---|---|---|---|---|---|---|---|---|
| Network | System | Network | Dice | $p_{Bonf}$ | Network | Dice | $p_{Bonf}$ | Network | Dice | $p_{Bonf}$ | Network | Dice | $p_{Bonf}$ | |
| PFN 1 | Default | Default B | 0.63 | 0.03 | Default | 0.41 | 0.02 | Default B | 0.52 | 0.02 | Default | 0.40 | 0.01 | Default Mode B |
| PFN 2 | SM | SM A | 0.68 | 0.07 | SM Foot | 0.88 | 0.02 | SM A | 0.64 | 0.06 | SM | 0.65 | 0.02 | Somatomotor Foot |
| PFN 3 | Frontopar | Control A | 0.65 | 0.02 | Frontopar | 0.62 | 0.02 | Frontopar A | 0.64 | 0.02 | Frontopar | 0.63 | 0.01 | Frontoparietal A |
| PFN 4 | SM | SM B | 0.48 | 0.13 | SM Face | 0.84 | 0.03 | SM B | 0.67 | 0.02 | Lateral SM | 0.83 | 0.01 | Somatomotor Face |
| PFN 5 | Dorsal | Dorsal A | 0.70 | 0.02 | Dorsal | 0.57 | 0.02 | Dorsal B | 0.76 | 0.02 | Dorsal | 0.44 | 0.02 | Posterior Dorsal Attention |
| PFN 6 | Visual | Visual B | 0.92 | 0.02 | Visual Med | 0.61 | 0.05 | Visual Peri | 0.93 | 0.02 | Visual | 0.62 | 0.02 | Visual Peripheral |
| PFN 7 | Ventral | Sal/Ventral B | 0.42 | 0.03 | Salience | 0.70 | 0.02 | Sal/ParMem | 0.49 | 0.02 | Salience | 0.34 | 0.01 | Salience |
| PFN 8 | Default | Temp Par | 0.69 | 0.03 | Lang | 0.68 | 0.02 | Lang | 0.52 | 0.02 | Ventral | 0.67 | 0.01 | Language |
| PFN 9 | Ventral | Sal/Ventral A | 0.67 | 0.02 | CingOp | 0.73 | 0.02 | CingOp | 0.69 | 0.02 | CingOp | 0.74 | 0.01 | Cingulo-Opercular |
| PFN 10 | Visual | Visual A | 0.67 | 0.07 | Visual Lat | 0.47 | 0.14 | Visual Cent | 0.87 | 0.02 | Visual | 0.41 | 0.19 | Visual Central |
| PFN 11 | SM | SM A | 0.31 | 0.44 | SM Hand | 0.47 | 0.02 | Premotor-PPr | 0.30 | 0.02 | SM | 0.32 | 0.22 | Somatomotor Right Hand |
| PFN 12 | Default | Default A | 0.50 | 0.02 | Default | 0.41 | 0.02 | Default A | 0.65 | 0.02 | Default | 0.41 | 0.01 | Default Mode A |
| PFN 13 | SM | SM A | 0.33 | 0.46 | SM Hand | 0.49 | 0.02 | Premotor-PPr | 0.28 | 0.03 | SM | 0.35 | 0.19 | Somatomotor Left Hand |
| PFN 14 | Dorsal | Dorsal B | 0.41 | 0.02 | Dorsal | 0.28 | 0.31 | Dorsal A | 0.35 | 0.02 | ParOcc | 0.23 | 0.04 | Anterior Dorsal Attention |
| PFN 15 | Frontopar | Control C | 0.48 | 0.03 | ParMem | 0.79 | 0.02 | Sal/ParMem | 0.27 | 0.03 | Med Parietal | 0.71 | 0.01 | Parietal Memory |
| PFN 16 | Auditory | SM B | 0.72 | 0.02 | Auditory | 0.84 | 0.02 | Auditory | 0.57 | 0.12 | Auditory | 0.80 | 0.01 | Auditory |
| PFN 17 | Frontopar | Control B | 0.54 | 0.02 | Default | 0.28 | 0.02 | Frontopar B | 0.51 | 0.02 | Frontopar | 0.28 | 0.04 | Frontoparietal B |

***Supplementary Table 6. Functional network labeling based on spatial correspondence of group atlas with 4 previously defined functional network atlases.*** To facilitate functional interpretation of our network results, we sought to give functional names to the 17 networks defined by our group atlas prior by testing spatial correspondence with 4 previously defined functional network atlases using the Network Correspondence Toolbox[18]. Specifically, the toolbox was implemented to calculate the Dice overlap between the hard parcellated group atlas with the Yeo-17[5], Gordon-17[19], Du-15[20], and Laumann-12[21] atlases, which uses spin testing to assess significance. Shown in the leftmost columns are the original network number labels and systems used in our prior work[1,2,22,23]. Subsequent columns show the best matching network in each atlas, defined as the highest Dice coefficient that passed the significance threshold of $p_{Bonf} < 0.05$ corrected across the number of comparisons. Highlighted cells indicate significant corresponding networks with functional labels with meaningful deviation from original Cui-17 system labeling, white unformatted cells indicate significant corresponding networks with labels consistent with original system labeling, and grayed-out cells indicate highest Dice overlap network in cases in which no network pairs reached significant overlap. Rightmost column shows final renamed network labels, informed by the most common label across the atlases with significant network matches, and in some cases further informed by cortical location of networks on the cortical surface (e.g., posterior vs. anterior dorsal attention, left vs. right hand networks). Notably, we kept system-level labeling consistent, with the understanding that networks operate across multiple scales[24] and thus used the 7 systems as a broader framework of functional organization. Network abbreviations: Cent = central; CingOp = cingulo-opercular; Dorsal: dorsal attention; Frontopar = frontoparietal; Lang = language; Lat = lateral; Med = medial; ParMem = parietal memory; ParOcc = parietal occipital; Peri = peripheral; PPr = posterior parietal rostral; Sal = salience; SM = somatomotor; Temp Par = temporal parietal; Ventral = ventral attention.

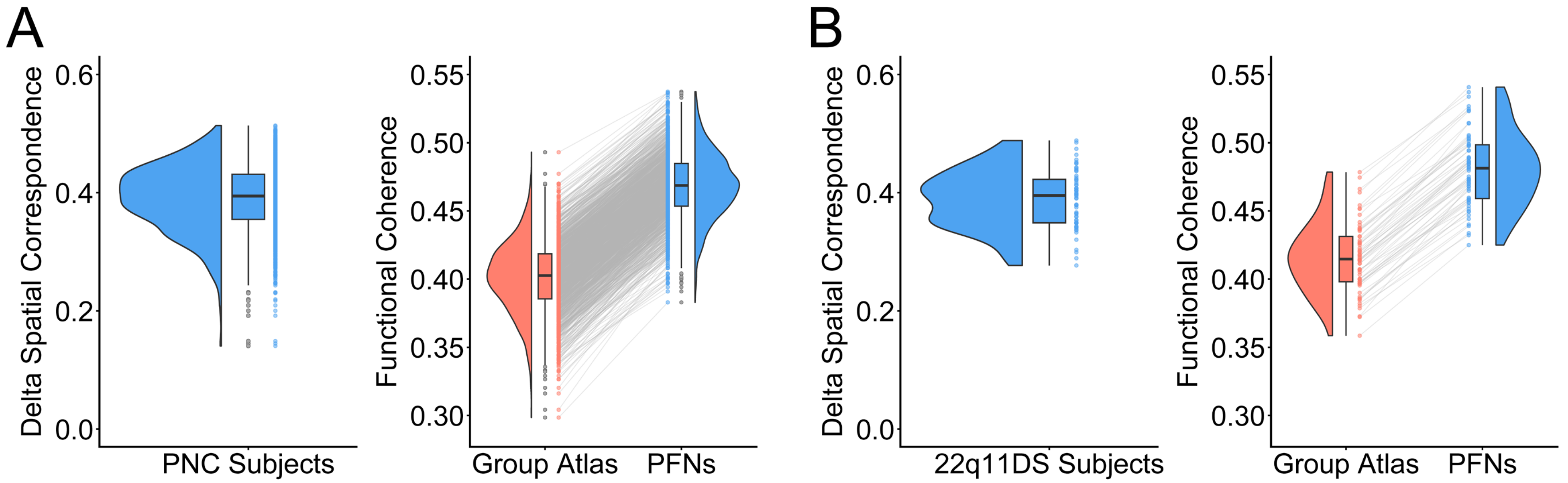


***Supplementary Figure 10. Personalized functional networks (PFNs) pass quality control measures of delta spatial correspondence and functional coherence in PNC and 22q11DS subjects.*** Personalized functional networks passed quality control measures utilized by the pNet toolbox[25] in both typically developing Philadelphia Neurodevelopmental Cohort (PNC) subjects (n=1332; **A**) as well as the matched 22q11.2 deletion syndrome (22q11DS) subjects (n=62; **B**). Delta spatial correspondence (left panels) tested whether the personalized network definition showed the greatest spatial correspondence with the homologous group atlas network, with a value greater than zero indicative of passing (correlation with homologous group atlas network minus strongest correlation with any nonhomologous group atlas network); plots show the lowest delta spatial correspondence across the 17 networks, with one data point per subject. Functional coherence (right panels) was defined as the loading-weighted mean of correlation coefficients between each vertex's time course and the network's loading-weighted mean time course. If our precision functional mapping method is capturing subject-specific functional organization more accurately, personalized network definitions would be expected to increase functional coherence relative to the group atlas. The plots show that averaged functional coherence across the 17 networks was increased in PFNs compared to group atlas definitions, with values paired at the subject-level. Boxplots represent the median and interquartile range.
