## Extended Data for "Personalized functional networks scale with brain size beyond anatomy and are reorganized in 22q11.2 deletion syndrome"

| | Total Vertices | Sig. Vertices | Sig. Vertices Prop. | Sig. Vertices Prop. Pos - Neg | Mean Beta of Sig. Vertices |
|---|---|---|---|---|---|
| VS-peri | 7,825 | 1,731 | 22.10% | −5.70% | −0.067 |
| VS-cent | 5,232 | 1,359 | 26.00% | 8.30% | 0.035 |
| **SM-foot** | 8,748 | 2,007 | 22.90% | −3.20% | −0.013 |
| SM-face | 4,975 | 1,476 | 29.70% | 0.40% | −0.024 |
| SM-rh | 6,971 | 2,211 | 31.70% | 5.70% | −0.002 |
| SM-lh | 5,537 | 923 | 16.70% | −9.10% | −0.120 |
| Auditory | 17,054 | 3,836 | 22.50% | 0.90% | −0.012 |
| DA-post | 18,222 | 5,424 | 29.80% | 0.40% | 0.018 |
| **DA-ant** | 16,764 | 6,501 | 38.80% | 21.20% | 0.082 |
| Salience | 12,838 | 4,474 | 34.80% | −0.10% | −0.031 |
| **CingOp** | 26,393 | 10,220 | 38.70% | −13.00% | −0.057 |
| FP-A | 24,726 | 7,985 | 32.30% | 2.80% | 0.009 |
| **FP-B** | 17,104 | 4,094 | 23.90% | 7.80% | 0.046 |
| **ParMem** | 4,504 | 1,117 | 24.80% | 13.00% | 0.089 |
| **Language** | 14,589 | 4,402 | 30.20% | −9.40% | −0.058 |
| **DM-B** | 21,215 | 7,141 | 33.70% | 12.40% | 0.052 |
| DM-A | 22,658 | 6,376 | 28.10% | −3.10% | −0.010 |
| | **235,355** | **71,277** | **30.30%** | | |

***Extended Data Table 1. Summary of vertex-level PFN topography changes with normative scaling.*** Bolded, colored networks correspond to those with significant deviations from isometry in functional areal scaling (Fig 2B, red = hyperallometric, blue = hypoallometric). Multiple comparisons in determining significant vertices corrected for using false discovery rate ($p_{FDR} < 0.05$). Vertex-level loading association directionality aligned with network-level functional areal results, as indicated by proportion of positive vs. negative significant vertices as well as mean β of significant vertices. Sig. = significant; Prop. = proportion. **Network abbreviations**: CingOp = cingulo-opercular; DA-ant = anterior dorsal attention; DA-post = posterior dorsal attention; ParMem = parietal memory; SM-lh = left hand; SM-rh = right hand; VS-cent = central visual; VS-peri = peripheral visual.

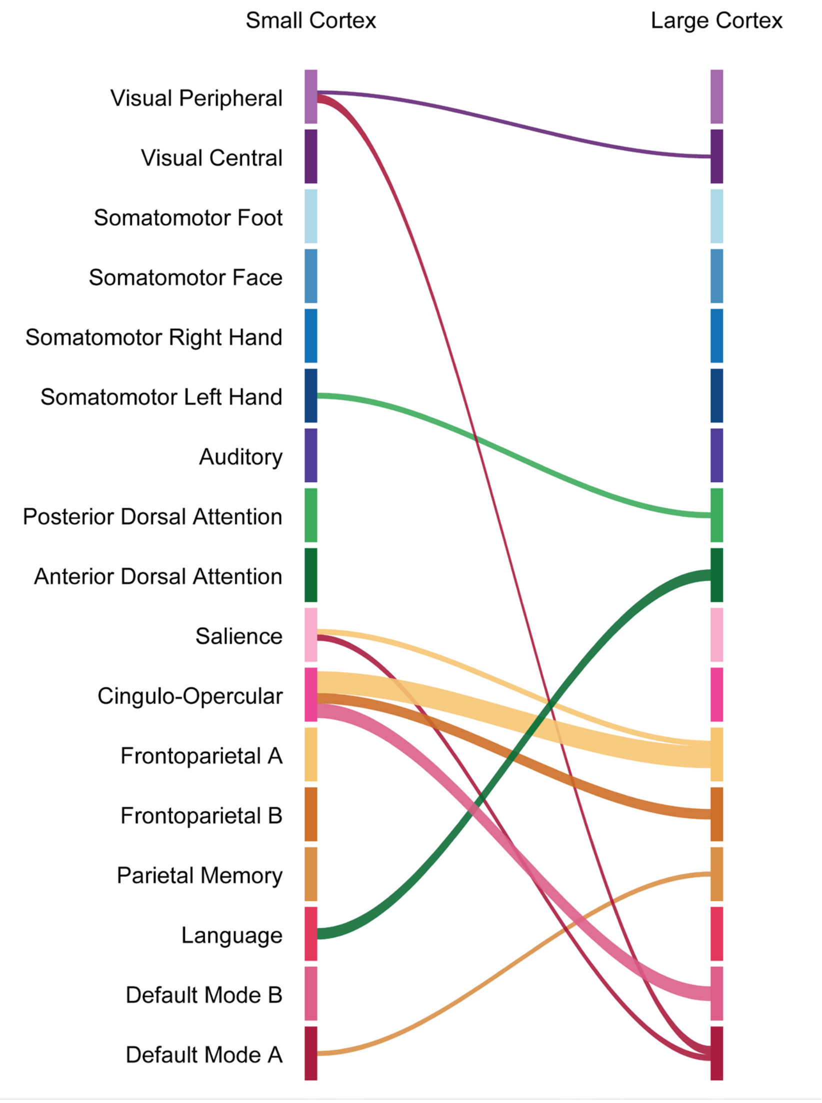


***Extended Data Figure 1. Network tradeoffs with increasing total cortical surface area in normative development.*** Sankey diagram displaying network territory tradeoffs with increasing total cortical surface area, as determined by significant FDR-thresholded directional reference masks for each PFN, defined as significant vertices whose β values matched the network's dominant direction. Dominant direction was determined by mean β across all significant vertices for each network; networks were only tested for tradeoff partners if their significant vertices in that direction ≥ 25 and absolute mean β across significant vertices ≥ 0.025. Tradeoff partners were identified as networks with the strongest opposite-direction averaged β within reference mask vertices (networks within 90% of the strongest tradeoff magnitude were retained). Ribbon color indicates network gaining territory with total cortical surface area; ribbon width reflects aggregate tradeoff burden, defined as the absolute product of the selected tradeoff network's mean β and the number of overlapping vertices. Network segments are shown at equal height for visualization and do not reflect network size. **Network abbreviations**: Aud = auditory; CingOp = cingulo-opercular; DA-ant = anterior dorsal attention; DA-post = posterior dorsal attention; DM = default mode; FP = frontoparietal; Lang = language; ParM = parietal memory; Sal = salience; SM = somatomotor; SM-lh = left hand; SM-rh = right hand; VS-cent = central visual; VS-peri = peripheral visual.

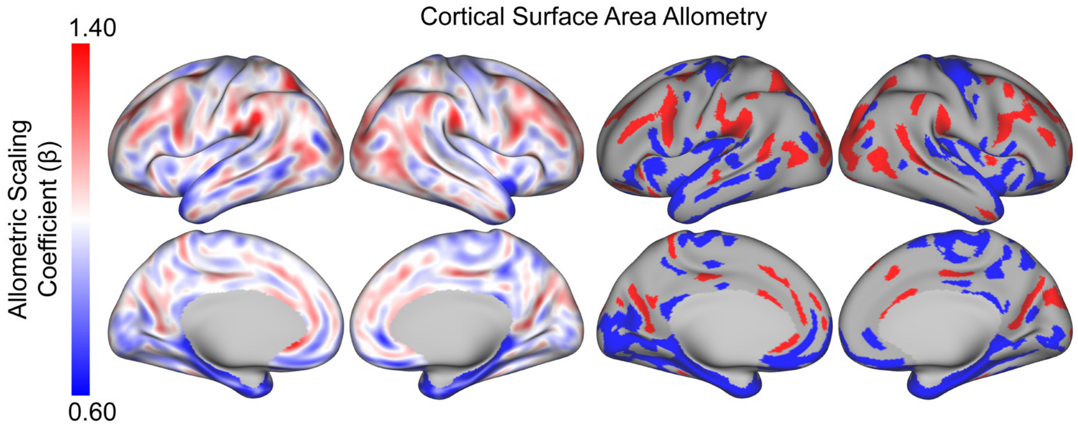


***Extended Data Figure 2. Normative anatomical surface area allometry.*** Normative anatomical allometry at the vertex-level, unthresholded (**left**). Map was significantly spatially correlated to prior results[1] in the same sample (PNC): r = 0.50, $p_{spin}$=2×10$^{-4}$. Significant effects ($p_{FDR}$ < 0.05) with 50 mm$^2$ cluster threshold (**right**).

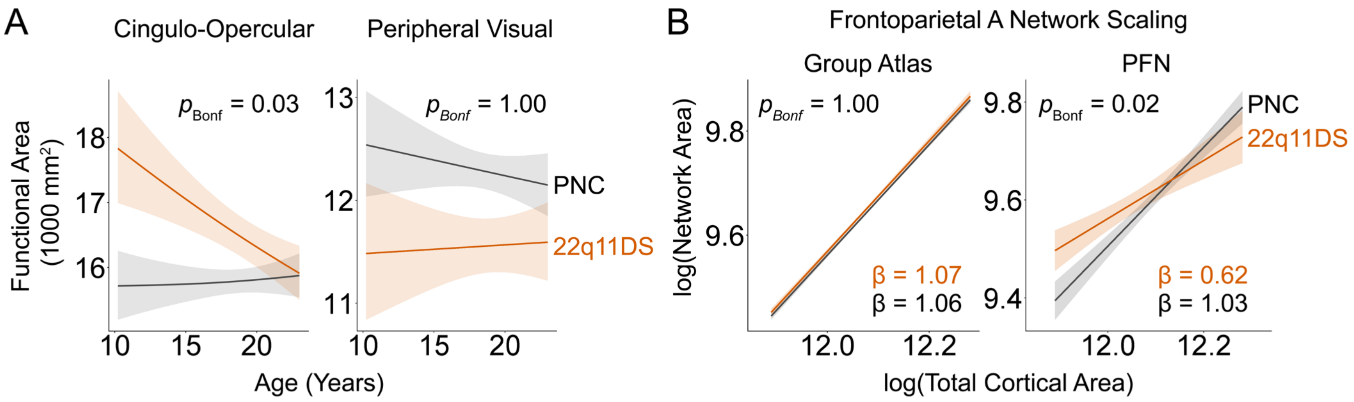


***Extended Data Figure 3. Significant interaction effects found between 22q11DS genotype with age and scaling using PFNs.*** **A.** A significant age-by-genotype interaction effect was found in the cingulo-opercular functional area, with attenuation of relative areal enlargement from ages 10 to 23 (common observed age range across cases and controls). In contrast, there was no significant age-by-genotype interaction effect observed in the peripheral visual network, which appears to remain disproportionately smaller in 22q11DS across this age range. Significance values correspond to interaction effects after Bonferroni correction. Curves represent population-standardized predictions of a generalized additive model (GAM) across the common observed age range based on an age spline, adjusted for fixed model covariates (total cortical area, sex, Euler number, and framewise displacement); predictions were obtained in log-space, and then back-transformed to the original area scale. **B.** Scaling-by-genotype interaction models revealed differential allometry in the frontoparietal A network depending on deletion status, an effect only detected in PFN-defined networks (right) but not in group atlas-defined networks (left). In the context of 22q11DS, frontoparietal A functional area appeared to display more strongly hypoallometric scaling (β=0.62) compared to allometry in matched PNC controls (β=1.03). Log-log plots represent population-standardized GAM predictions across the common observed total cortical area range, adjusted for fixed model covariates (age spline, sex, Euler number, and framewise displacement); predictions remain in log-space.

| | Scaling Unadjusted | | | | | Scaling Adjusted | | | |
|---|---|---|---|---|---|---|---|---|---|
| | Total Vertices | Sig. Vertices | Sig. Vertices Prop. | Sig. Vertices Prop. Pos - Neg | Mean Beta of Sig. Vertices | Sig. Vertices | Sig. Vertices Prop. | Sig. Vertices Prop. Pos - Neg | Mean Beta of Sig. Vertices |
| **VS-peri** | 7,825 | 42 | 0.54% | −0.54% | −0.083 | 62 | 0.79% | −0.77% | −0.089 |
| VS-cent | 5,232 | 20 | 0.38% | 0.38% | 0.081 | 28 | 0.54% | 0.54% | 0.075 |
| SM-foot | 8,748 | 118 | 1.35% | 1.33% | 0.053 | 76 | 0.87% | 0.87% | 0.060 |
| SM-face | 4,975 | 97 | 1.95% | −0.10% | 0.005 | 40 | 0.80% | −0.24% | −0.024 |
| SM-rh | 6,971 | 232 | 3.33% | 0.49% | −0.001 | 152 | 2.18% | 1.32% | 0.037 |
| SM-lh | 5,537 | 82 | 1.48% | 1.48% | 0.086 | 15 | 0.27% | 0.27% | 0.066 |
| Auditory | 17,054 | 114 | 0.67% | −0.08% | −0.017 | 55 | 0.32% | −0.09% | −0.027 |
| DA-post | 18,222 | 319 | 1.75% | 1.53% | 0.074 | 211 | 1.16% | 1.16% | 0.087 |
| DA-ant | 16,764 | 271 | 1.62% | −1.53% | −0.079 | 181 | 1.08% | −0.97% | −0.072 |
| Salience | 12,838 | 377 | 2.94% | 0.35% | −0.007 | 184 | 1.43% | 0.37% | 0.010 |
| **CingOp** | 26,393 | 460 | 1.74% | 1.44% | 0.074 | 268 | 1.02% | 1.01% | 0.093 |
| FP-A | 24,726 | 395 | 1.60% | 0.17% | 0.006 | 177 | 0.72% | −0.37% | −0.039 |
| FP-B | 17,104 | 189 | 1.11% | 0.66% | 0.036 | 55 | 0.32% | 0.17% | 0.029 |
| ParMem | 4,504 | 94 | 2.09% | −1.51% | −0.062 | 6 | 0.13% | 0.00% | −0.008 |
| Language | 14,589 | 15 | 0.10% | 0.01% | 0.009 | 8 | 0.05% | −0.05% | −0.044 |
| DM-B | 21,215 | 351 | 1.65% | −0.35% | −0.019 | 189 | 0.89% | −0.01% | −0.006 |
| DM-A | 22,658 | 159 | 0.70% | 0.57% | 0.055 | 101 | 0.45% | 0.30% | 0.046 |
| | 235,355 | **3,335** | **1.42%** | | | **1,808** | **0.77%** | | |

***Extended Data Table 2. Summary of vertex-level PFN topography changes with 22q11DS genotype.*** Bolded, colored networks in first column correspond to significant 22q11DS functional areal effects (Fig 4C right, red = positive, blue = negative). Multiple comparisons in determining significant vertices corrected for using false discovery rate ($p_{FDR} < 0.05$). Vertex-level loading association directionality aligned with significant network-level functional areal results, as indicated by proportion of positive vs. negative significant vertices as well as mean β of significant vertices. Directional pattern of significant loading changes across networks remained broadly consistent across scaling unadjusted and adjusted models, with attenuation in number of significant vertices. Sig. = significant; Prop. = proportion. **Network abbreviations**: CingOp = cingulo-opercular; DA-ant = anterior dorsal attention; DA-post = posterior dorsal attention; ParMem = parietal memory; SM-lh = left hand; SM-rh = right hand; VS-cent = central visual; VS-peri = peripheral visual.

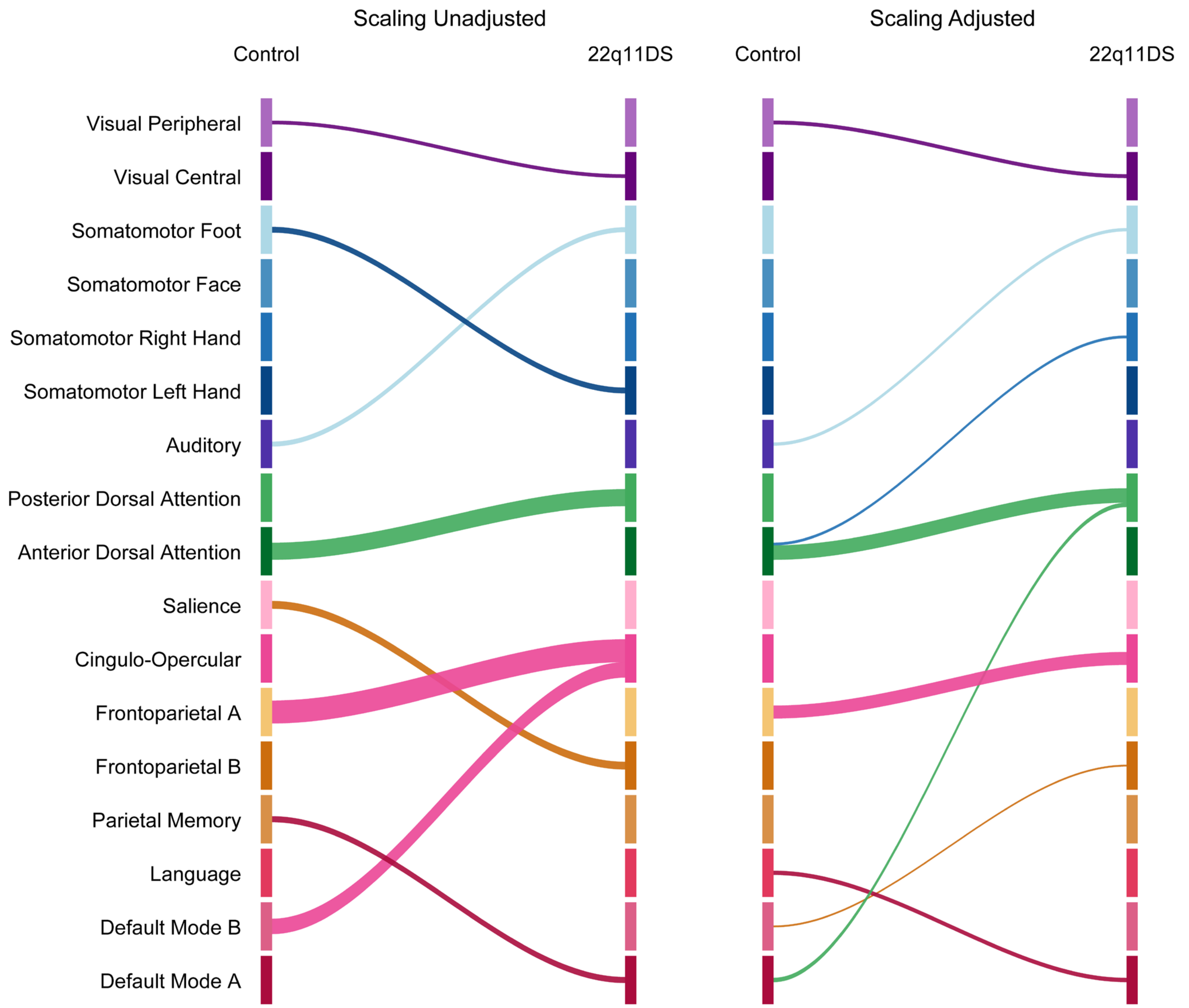


***Extended Data Figure 4. Network tradeoffs with 22q11DS, before and after scaling adjustment.*** Sankey diagrams displaying network territory tradeoffs associated with 22q11DS, before (**A**) and after total cortical surface area adjustment (**B**). Tradeoff derivation described in Extended Data Figure 1. Ribbon width is magnified 25-fold compared to Extended Data Figure 1 to improve visibility. Network tradeoff patterns remained broadly consistent before and after scaling adjustment; however, tradeoffs do appear attenuated after scaling adjustment. **Network abbreviations**: Aud = auditory; CingOp = cingulo-opercular; DA-ant = anterior dorsal attention; DA-post = posterior dorsal attention; DM = default mode; FP = frontoparietal; Lang = language; ParM = parietal memory; Sal = salience; SM = somatomotor; SM-lh = left hand; SM-rh = right hand; VS-cent = central visual; VS-peri = peripheral visual.

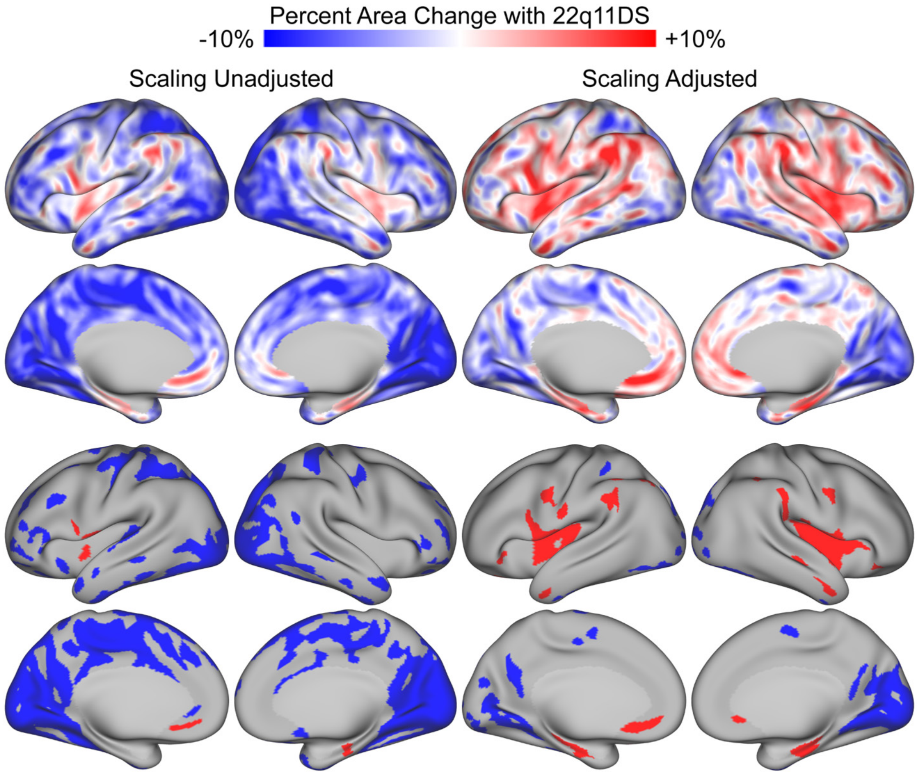


***Extended Data Figure 5. Cortical surface area changes with 22q11DS.*** Anatomical surface area effects of 22q11DS at the vertex-level, before and after scaling adjustment (**left** and **right**, respectively). Significant effects ($p_{FDR} < 0.05$) with 50 mm$^2$ cluster threshold shown in bottom row.
